# Polygenic risk for schizophrenia converges on alternative polyadenylation as molecular mechanism underlying synaptic impairment

**DOI:** 10.1101/2024.01.09.574815

**Authors:** Florian J. Raabe, Anna Hausruckinger, Miriam Gagliardi, Ruhel Ahmad, Valeria Almeida, Sabrina Galinski, Anke Hoffmann, Liesa Weigert, Christine K. Rummel, Vanessa Murek, Lucia Trastulla, Laura Jimenez-Barron, Alessia Atella, Susanne Maidl, Danusa Menegaz, Barbara Hauger, Eva-Maria Wagner, Nadia Gabellini, Beate Kauschat, Sara Riccardo, Marcella Cesana, Sergi Papiol, Vincenza Sportelli, Monika Rex-Haffner, Sebastian J. Stolte, Michael C. Wehr, Tatiana Oviedo Salcedo, Irina Papazova, Sevilla Detera-Wadleigh, Francis J McMahon, Andrea Schmitt, Peter Falkai, Alkomiet Hasan, Davide Cacchiarelli, Udo Dannlowski, Igor Nenadić, Tilo Kircher, Volker Scheuss, Matthias Eder, Elisabeth B. Binder, Dietmar Spengler, Moritz J. Rossner, Michael J. Ziller

## Abstract

Schizophrenia (SCZ) is a genetically heterogenous psychiatric disorder of highly polygenic nature. Correlative evidence from genetic studies indicate that the aggregated effects of distinct genetic risk factor combinations found in each patient converge onto common molecular mechanisms. To prove this on a functional level, we employed a reductionistic cellular model system for polygenic risk by differentiating induced pluripotent stem cells (iPSCs) from 104 individuals with high polygenic risk load and controls into cortical glutamatergic neurons (iNs). Multi-omics profiling identified widespread differences in alternative polyadenylation (APA) in the 3’ untranslated region of many synaptic transcripts between iNs from SCZ patients and healthy donors. On the cellular level, 3’APA was associated with a reduction in synaptic density of iNs. Importantly, differential APA was largely conserved between postmortem human prefrontal cortex from SCZ patients and healthy donors, and strongly enriched for transcripts related to synapse biology. 3’APA was highly correlated with SCZ polygenic risk and affected genes were significantly enriched for SCZ associated common genetic variation. Integrative functional genomic analysis identified the RNA binding protein and SCZ GWAS risk gene PTBP2 as a critical trans-acting factor mediating 3’APA of synaptic genes in SCZ subjects. Functional characterization of PTBP2 in iNs confirmed its key role in 3’APA of synaptic transcripts and regulation of synapse density. Jointly, our findings show that the aggregated effects of polygenic risk converge on 3’APA as one common molecular mechanism that underlies synaptic impairments in SCZ.

## Introduction

Schizophrenia (SCZ) is a highly debilitating psychiatric disorder originating from the complex interplay of genetic and environmental factors^1^. Genetics has offered a compelling handle on the biological basis of SCZ due its overall high heritability of 79%^2^. Genome Wide Association Studies (GWAS) identified more than 270 common^3^ and 32 rare genetic risk factors^4^, underscoring the highly polygenic architecture of disease risk in SCZ. Similar results have been obtained for other psychiatric disorders such as major depression (MDD)^5^ and bipolar disorder (BD)^6^. Genomic and pathway level annotation studies based on the biological role of the genes within these risk loci implicated synapse, immune and transcriptional control as key processes altered in SCZ and other psychiatric diseases^7,8^. Moreover, scRNAseq from post-mortem brains showed that the expression of SCZ risk genes was highly enriched in excitatory and inhibitory neurons of the cortex^3,4^.

However, the molecular mechanisms underlying the de-regulation of these genes in disease as well as their contribution to disease etiology at the functional level remain largely unknown.

To address the former challenge, previous studies focused on the functional characterization of individual common variants (CVs) and the associated genes identified in GWA studies^9,10^. These analyses revealed how disease associated variants alter the regulation of individual genes in *cis*, including modulation of gene expression levels, the chromatin state of regulatory elements or alterations of gene splicing patterns, effectively acting as quantitative trait loci (QTL)^11–13^. Jointly, these molecular mechanisms are currently estimated to modulate 20-30% of disease associated GWAS loci^11,12,14^, indicating the presence of additional yet unknown molecular mechanisms contributing to impairment of gene function.

Such variant and gene-centric studies greatly improved our understanding of the biological function of individual genetic variants. However, their impact on our understanding how these individual variant effects converge on common molecular mechanisms contributing to the etiology of mental illness remains limited.

This limitation is rooted in the high prevalence of common disease associated genetic variants in the general population. Therefore, each variant individually contributes only minimally to overall disease risk and is in isolation not capable to drive disease relevant alterations in molecular and cellular processes. Instead, its disease relevant biological function is highly dependent on the presence of other genetic risk variants. Moreover, due to the highly polygenic architecture of disease risk in SCZ, the patient population is characterized by an extreme heterogeneity in genetic risk variant distribution, leaving almost every patient with an individual genetic risk factor profile. Thus, in a complementary line of research, large cohort studies sought to pinpoint the aggregated effects of genetic risk factors as well as life and treatment history on postmortem brain samples from individuals with psychiatric disorders and neurotypical controls^11,13^. Although these analyses identified widespread changes in gene expression and splicing patterns in the prefrontal cortex (PFC) of adult individuals with SCZ, BD or ASD, associations between transcriptional changes with common genetic risk factors and in particular polygenic risk were modest to small. The latter observation likely reflects a strong impact of a lifelong disease history on the transcriptome as well as the relevance of additional molecular mechanisms in mediating the converging effect of polygenic risk factors in mental illness. In this respect, post-transcriptional mRNA regulation constitutes an additional disease relevant molecular mechanism potentially mediating polygenic risk factor effects, given its prominent role in neuronal physiology^15^ and complex genetic regulation^16,17^.

Here, we sought to translate the effects of highly heterogenous polygenic risk across individuals into common molecular mechanisms underlying the etiology in SCZ. Therefore, we set out to pinpoint the joint functional effects of genetic risk factors on a wide array of molecular and cellular phenotypes in a genetic model system relevant to mental illness.

Therefore, we leveraged the unique feature of iPSCs to capture the polygenetic risk architecture of each individual and assessed the aggregated functional consequences of these genetic risk factors on molecular and cellular endophenotypes in the absence of patient level environmental confounders.

Consistent with previous studies, these analyses identified moderate changes in the transcriptome and epigenome of neuronal cells from individuals with SCZ (ISCZ) and the absence of association with polygenic risk. In stark contrast, analysis of post-transcriptional alterations revealed that highly heterogeneous polygenic risk converges on the alternative polyadenylation of the 3’ untranslated region (3’APA) of many key synaptic genes. Moreover, we identify de-regulation of the trans-acting RNA binding protein and SCZ risk gene *PTBP2* as one critical contributor to 3’APA in SCZ and demonstrate its role in modulating synaptic density as disease relevant cellular endophenotype.

## A comprehensive iPSC cohort from patients with mental illness

We assembled a large cohort of individuals from European ancestry with major psychiatric disorders as well as healthy controls from several research cohorts^18–21^ (**Fig. 1a**). Given that SCZ genetically correlates with bipolar disorder (BD) and unipolar major depressive disorder (MDD)^22,23^, we also included individuals with these diseases to evaluate the specificity of polygenically driven effects. This cohort (n=104) comprised neurotypical (healthy) controls (HC, n = 38), and individuals with SCZ (ISCZ, n=38), BD (n = 20) and MDD (n = 8) (**Supplementary Table 1**) with a mean age of onset in the late 20s adolescence (mean 28.5/27.1/29.1 SCZ/BD/MDD).

**Fig. 1.**
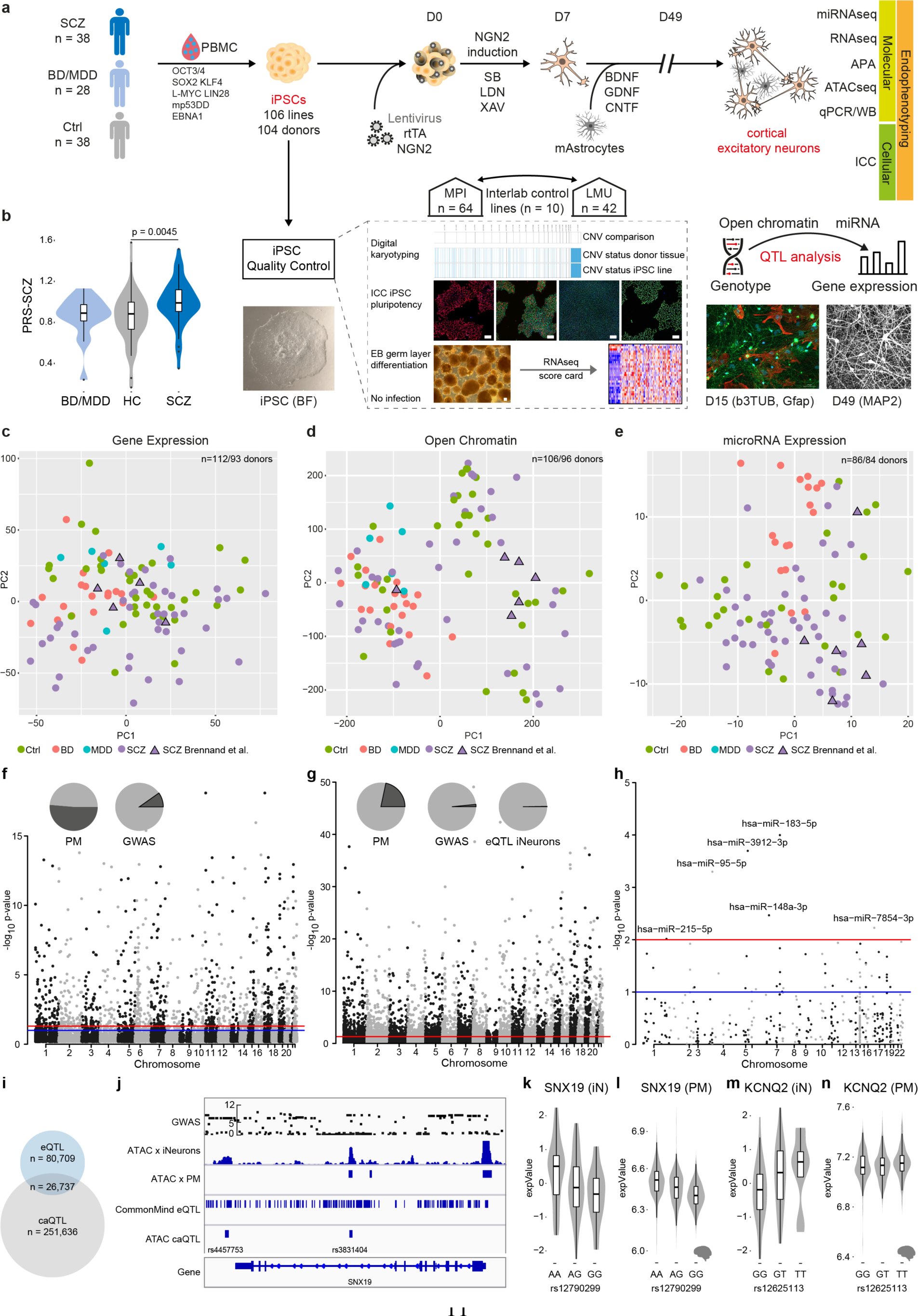
Multi-omic profiling of iNeuron mental health cohort and quantitative trait analysis for gene expression, open chromatin and microRNA expression. **a**, Schematic overview of iPSC line generation and differentiation to NGN2-directed cortical excitatory neurons (iNs), co-culturing with murine Astrocytes, and multi-modal endophenotyping after 49 days of differentiation which includes RNAseq, ATACseq, miRNAseq, qPCR, Western Blot (WB) on the molecular level, immunocytochemistry (ICC), fluorescence in situ hybridization (FISH) and electrophysiology by patch-clamp on a cellular level as well as multielectrode arrays (MEA) on the (micro)circuit level in a multicenter approach in order to perform a case-control study with patients affected with schizophrenia (SCZ), unipolar major depressive disorder (MDD), bipolar disorder (BD) and healthy controls (HC) to perform QTL analysis. iPSC characterization pipeline includes digital karyotyping using microarrays, immunocytochemistry (ICC) analysis of key pluripotency markers, and assessment of differentiation potential by embryoid body (EB) generation followed by RNA-Seq to perform score card assessment of the expressed germ layer signature. Infections with HIV, HCV, CMV and mycoplasma were excluded. Scale bars indicate 50µm. **b**, Distribution of the individual polygenic risk score for schizophrenia (PGC3 SCZ) for the different diagnosis groups of hiPSC donors. P-value based on two-sided Wilcoxon-Test. **c**, Principal component analysis of RNA-Seq based gene expression profiles across iN samples from 93 distinct donors at day 49. Different colors indicate diagnosis group and triangles highlight iPSC line donors originally employed by Brennand and colleagues and subsequently used in many investigations in the field for reference. The latter were re-derived from the same families for this study. **d**, Principal component analysis of ATAC-Seq based open chromatin profiles across iN samples from 96 distinct donors at day 49. Different colors indicate diagnosis group and triangles highlight iPSC line donors originally employed by Brennand and colleagues for reference. **e**, Principal component analysis of microRNA-Seq based profiles across 86 iNs samples from 86 distinct donors at day 49. Different colors indicate diagnosis group and triangles highlight iPSC line donors originally employed by Brennand and colleagues for reference. **f**, Manhatten plot of eQTL association strength (y-axis) across all chromosomes (x-axis). Red line indicates global significance level after multiple testing correction, identifying 1,331 genes with at least one eQTL (FDR:<0.05). Pie charts indicate overlap (dark grey) of identified SNPs with eQTLs found in human post mortem DLPFC (left) or overlapping with SCZ GWAS risk loci (right). **g**, Manhatten plot of caQTL association strength (y-axis) across all chromosomes (x-axis). Red line indicates global significance level after multiple testing correction, identifying 4,904 peaks with at least one caQTL (Bonferroni corrected p-value :< 0.01). Pie charts indicate overlap (dark grey) of identified SNPs with eQTLs found in human post mortem DLPFC (left), overlapping with SCZ GWAS risk loci (middle) or with eQTLs identified in iNs (right). **h**, Manhatten plot of mirQTL association strength (y-axis) across all chromosomes (x-axis). Red line indicates global significance of permutation based p-value, identifying 6 microRNAs (p-value :< 0.01) with at least one mirQTL. **i**, Venn diagram showing the overlap of SNPs identified as eQTL and caQTL associated SNPs. **j**, Representative example of genomic annotations and distinct QTL types at the sorting nexin 19 (*SNX19)* locus, left: from top to bottom: Association strength of imputed SNPs at the locus with SCZ based on SCZ GWAS from Ripke et al. 2013, ATAC-Seq profile of representative iNs at day 49 and post mortem cortex, eQTLs detected at the locus in iNs, caQTLs detected at the locus co-localizing with open chromatin regions. **k**, Distribution of normalized gene expression level of SNX19 in iNs (y-axis) by allele status (x-axis). **l**, Distribution of normalized gene expression level of SNX19 (y-axis) in human post mortem DLPFC by allele status (x-axis). **m**, Distribution of normalized gene expression level of the voltage gate potassium channel, subfamily Q, member 2 (*KCNQ2* y-axis) in iNs by allele status. **n**, Distribution of normalized gene expression level of *KCNQ2* (y-axis) in human post mortem DLPFC by allele status (x-axis).

Importantly, ISCZ exhibited increased polygenic risk scores for SCZ (PRS-SCZ) (p= 0.0045, two-sided Wilcoxon-Test; **Fig. 1b**) concomitant with the general absence of copy number variations previously associated with SCZ (**Extended Data Fig. 1a**). Moreover, all individuals were characterized by a highly heterogeneous distribution of genetic risk factors for SCZ based on common genetic variants identified by GWAS (**Extended Data Fig. 1b**), implying private genetic risk factor profiles. In contrast, no difference in PRS for BD or MDD was detected between any diagnosis group (**Extended Data Fig. 1c,d**). Finally, our cohort also encompasses a set of 4 ISCZ whose cells have been investigated in many pioneering/previous iPSC studies on SCZ as additional points of comparison^25^.

We generated 106 iPSC lines from the present cohort and previous set^25^ and subjected them to comprehensive quality control (**Fig. 1a**) including digital karyotyping (**Supplementary Data 1**), analysis of pluripotency (**Supplementary Data 2**), differentiation potential and lineage marker analysis (**Extended Data Fig. 1e-l**).

All iPSC lines passing quality control and devoid of genetic aberrations were differentiated into iNs over a timeline of 49 days, relying on lentiviral transduction of Neurogenin-2 (*Ngn2*) in conjunction with small molecule application^26^ (**Fig. 1a**). To facilitate cross-lab reproducibility, the iPSC cohort was split in two groups and differentiation experiments were performed at two distinct sites (**Fig. 1a**). For iN analysis, we established a comprehensive multi-layered endophenotype pipeline including deep molecular phenotyping by RNA-Seq and alternative polyadenylation analysis (n=93 donors), ATAC-Seq (n=96 donors) as well as subsets for microRNA-Seq (n=85 donors) followed by high content imaging (n=20 donors) for synaptic density assessment (**Fig. 1a**).

## Molecular characteristics of iNs from patients with psychiatric disorders

RNA-Seq and immunocytochemistry (ICC) based analysis confirmed differentiation into well-matured excitatory iNs (**Extended Data Fig. 2a-c**) in line with previous reports^26^. Single cell RNA-Seq (scRNA-Seq) analysis of iNs from 4 donors confirmed bulk and ICC measurements, revealing 7 distinct clusters, mostly differing by excitatory neuron subtype marker expression and/or maturation stage (**Extended Data Fig. 2d-f**). *In silico* deconvolution of iN bulk RNA-Seq profiles using marker gene sets derived from each cluster of the scRNA-Seq iN data^27^ revealed uniform differentiation across the lines with little variation in cellular composition for BD iNs and no difference between iNs from HC and ISCZ (**Extended Data Fig. 2g**). These results underscore the absence of gross differences in cellular identity and distribution or developmental trajectory among iNs from cases and controls. Similarly, basic electrophysiological properties were comparable between neurons from different diagnosis groups and comparable to mature primary human or mouse neurons^28^ except for differences in input resistance (**Extended Data Fig. 2h-i**). In line with these findings and iN marker expression (**Extended Data Fig. 2a**), iNs most closely cluster with excitatory neurons of progressed post-natal development (P7) (**Extended Data Fig. 2j**) based on previously published transcriptomic signatures of the developing mouse cortex^29^. Finally, principal component analyses of the RNA-Seq, ATAC-Seq and microRNA-Seq datasets revealed no clustering by diagnostic group in either of the modalities (**Fig. 1c-e**). Instead, diagnostic groups were highly interspersed, indicating no fundamental differences in cellular state between the diagnosis groups.

## QTL mapping across molecular features identifies multi-layered mechanisms of cis-acting genetic variation contributing to polygenic risk in SCZ

Gene expression, open chromatin and microRNA expression levels constitute a critical substrate of psychiatric disorder associated genetic variation^30,31^. Therefore, we sought to pinpoint *cis*-acting variants driving alterations in these molecular layers in iNs across patients and healthy donors. The quantitative trait locus (QTL) analysis identified 1,331 genes (eQTLs), 4,904 open chromatin peaks (caQTLs), 6 microRNAs (miQTLs) subject to modulation by genetic variants in *cis* (**Fig. 1f-h**). Importantly, 55.4% of the identified eQTLs overlapped with eQTLs identified in adult human tissue from the prefrontal cortex (PFC)^11^, supporting their *in vivo* relevance (**Fig. 1f** and **Supplementary Table 2**). For open chromatin and microRNAs, no large publicly available reference datasets from relevant primary tissues are currently available to confirm their overlap. Nevertheless, 21.7% of the identified open chromatin peaks associated with caQTLs overlapped with eQTLs detected in bulk PFC. (**Fig. 1g** and **Supplementary Table 2**).

Interestingly, the overlap of QTLs across molecular features in iNs was limited, with 24.9% of eQTL overlapping with caQTLs (**Fig. 1l** and **Supplementary Table 2**). These observations support the notion that distinct *cis* acting variations affect different molecular layers with likely distinct downstream effects. Moreover, 28 genes, 78 peaks and no microRNA with e/ca/miQTLs overlapped with GWAS risk loci for SCZ (**Fig. 1f-h** and **Supplementary Table 2**).

Among the identified QTLs co-localizing with SCZ GWAS risk loci was a caQTL at the *SNX19* and the *KCNQ2* locus (both also observed in PFC) (**Fig. 1j-n**) as well as multiple multiple caQTL in intronic regions of *NRXN1* (**Extended Data Fig. 3a**). This is of particular interest as one of these caQTLs is located in an alternative promoter of *NRXN1* and thus potentially contributes to the previously reported isoform abundance diversity of *NRXN1* in ISCZ^32^.

In summary, these observations underscore the capacity of iNs to capture a substantial fraction of the functional effects of genetic variation on disease relevant molecular layers from human PFC. Moreover, these findings also reveal a substantial diversity in the affected molecular features in *cis*, as distinct common variants modulate gene expression, microRNA expression or chromatin accessibility.

## Identification of differentially regulated molecular features in iNs from ISCZ

Going beyond individual *cis*-mediated genetic effects, we next leveraged the unique feature of iNs to assess the aggregated functional consequences of distinct polygenic risk profiles in SCZ (**Figures 1b**, **Extended Data Fig. 1a,b**) in the absence of medication and environmental confounders. To that end, we investigated an array of molecular layers in iNs from HC and ISCZ, including gene expression, microRNA expression, open chromatin marks and alternative polyadenylation as post-transcriptional regulatory mechanism.

Following a detailed power evaluation (**Extended Data Fig. 3b-d**), we performed differential expression analysis and identified 467 transcripts (365 protein coding and 102 non-coding transcripts) (FDR≤0.01, |log_2_FC|≥0.4; **Fig. 2a** and **Extended Data Fig. 3e**, and **Supplementary Table 3**), henceforth referred to as DEGs as well as 67 micro RNAs (FDR≤0.05; **Fig. 2b** and **Supplementary Table 3)**. For open chromatin regions, no peaks with differential accessibility were detected significant after multiple testing correction (FDR:<0.05; **Extended Data Fig. 3f**) despite high reproducibility of the generated profiles (**Extended Data Fig. 3g**). This might reflect overall lower sensitivity of the ATAC based data.

**Fig. 2.**
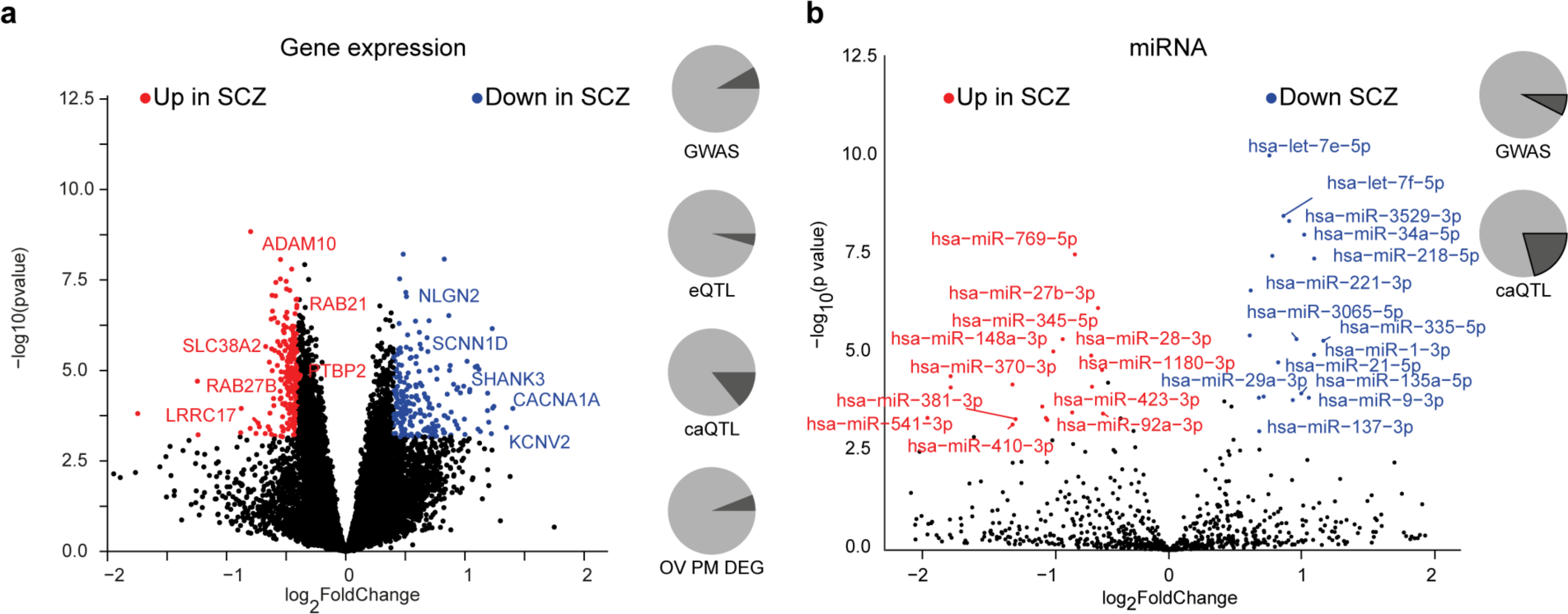
Differential molecular feature analysis between SCZ and HC iPSC-derived neurons. **a**, Volcano plot showing the log_2_ fold-change (x-axis) and significance (-log_10_ p-value, y-axis) of differentially expressed genes (DEGs) between SCZ and HC iNs at day 49. Positive fold-changes indicate lower expression in SCZ. Red/blue dots indicate significance (FDR:<0.01) and minimal fold-change (|fc|=ζ0.4) cutoffs to define DEGs. Text highlights selected genes differentially expressed. Pie charts from top to bottom show: overlap (dark grey) of DEGs with GWAS associated genes, iNs-based eQTLs, iNs-based caQTLs within 100kb and fraction of genes detected as also differentially expressed between HC and SCZ PFC postmortem (p-value :< 0.01). **b**, Volcano plot showing the log_2_ fold-change and significance of differential microRNA expression between SCZ and HC iNs at day 49. Positive fold-changes indicate lower expression in SCZ. Red/blue dots indicate significance (FDR:<0.01) and minimal fold-change (|fc|ζ0.5) cutoffs to define DE microRNAs. Text highlights selected microRNAs DE. Pie charts show overlap (dark grey) of DE microRNAs with GWAS associated genes and iNs-based caQTLs within 200kb.

Among the differentially expressed microRNAs, we identified several miRs that have been implicated previously in SCZ by GWAS (e.g. miR-135a, miR-137, miR-1180)^33^ and/or with their targetome connected to SCZ risk genes (e.g. miRNA-1, miR-7-5p, mir-9-5p)^34,35^. Moreover, several differentially expressed microRNAs in iNs were also found to be differentially present in peripheral blood of SCZ patients (e.g. let-7g, miR-7, miR-34a-5p, miR-218-5p, miR-132)^36,37^ (**Fig. 2b**), suggesting their trans-tissue and potentially diagnostic relevance.

The set of DEGs included 39 SCZ GWAS risk genes while 18 were associated with eQTLs and 36 with caQTL in iNs (**Fig. 2a**). Moreover, 36 DEGs overlapped with DEGs from human PFC comparing neurotypical controls and ISCZ (**Fig. 2a**). However, mean expression of DEGs across samples did not associate with SCZ-PRS (p=0.13, r2=-0.169). Lastly, we note a high disease specificity of DEGs in the SCZ (n=467) and the BD/MDD (n=11 genes) cohorts compared to the neurotypical group with only 1 gene (*PPIAP11*) being shared among the two psychiatric disorders (**Extended Data Fig. 3h**). Notably, we also detected expression differences in multiple genes critical for synaptic function and neuronal activity^38^ such as *NLGN2, SHANK3, RAB21, RAB27B, ADAM10, CACNA1A and SCNN1D* to be downregulated in iNs from ISCZ (**Fig. 2a**). Pathway level enrichment did not yield any significant results for genes up-regulated in ISCZ and implicated protein metabolism related processes for down-regulated genes. Consistent with previous reports^13^, most of the DEGs, including those with roles in synaptic and neuronal activity, do not harbor any association signal in SCZ or BD GWAS in *cis* range.

## Widespread 3’APA of synaptic genes in ISCZ

Going beyond previously investigated molecular traits, we next assessed post-transcriptional mRNA regulation as potential mechanism mediating polygenic risk effects in SCZ. We focused on alternative polyadenylation of the 3’ untranslated region (UTR) as the non-protein coding section of the messenger RNA that follows the stop codon. This region contains numerous RNA cis-regulatory elements relevant for the post-transcriptional regulation of the mRNA, modulating RNA stability, translation rates and subcellular localization^39^. Thus, alternative length and composition of the 3’UTR can have a profound impact on these properties of the mRNA, resulting in a critical role in neuronal physiology and synapse biology^15^.

We mapped polyadenylation sites (PAS) in iNs from ISCZ using 3’RNA-Seq and identified a total of 19,960 PAS of which 7,812 were not previously detected (**Fig. 3a,b**). Subsequently, we leveraged this library of PAS in combination with our iNs RNA-Seq dataset to identify differential 3’UTR APA events between HC and ISCZ^40^.

**Fig. 3:**
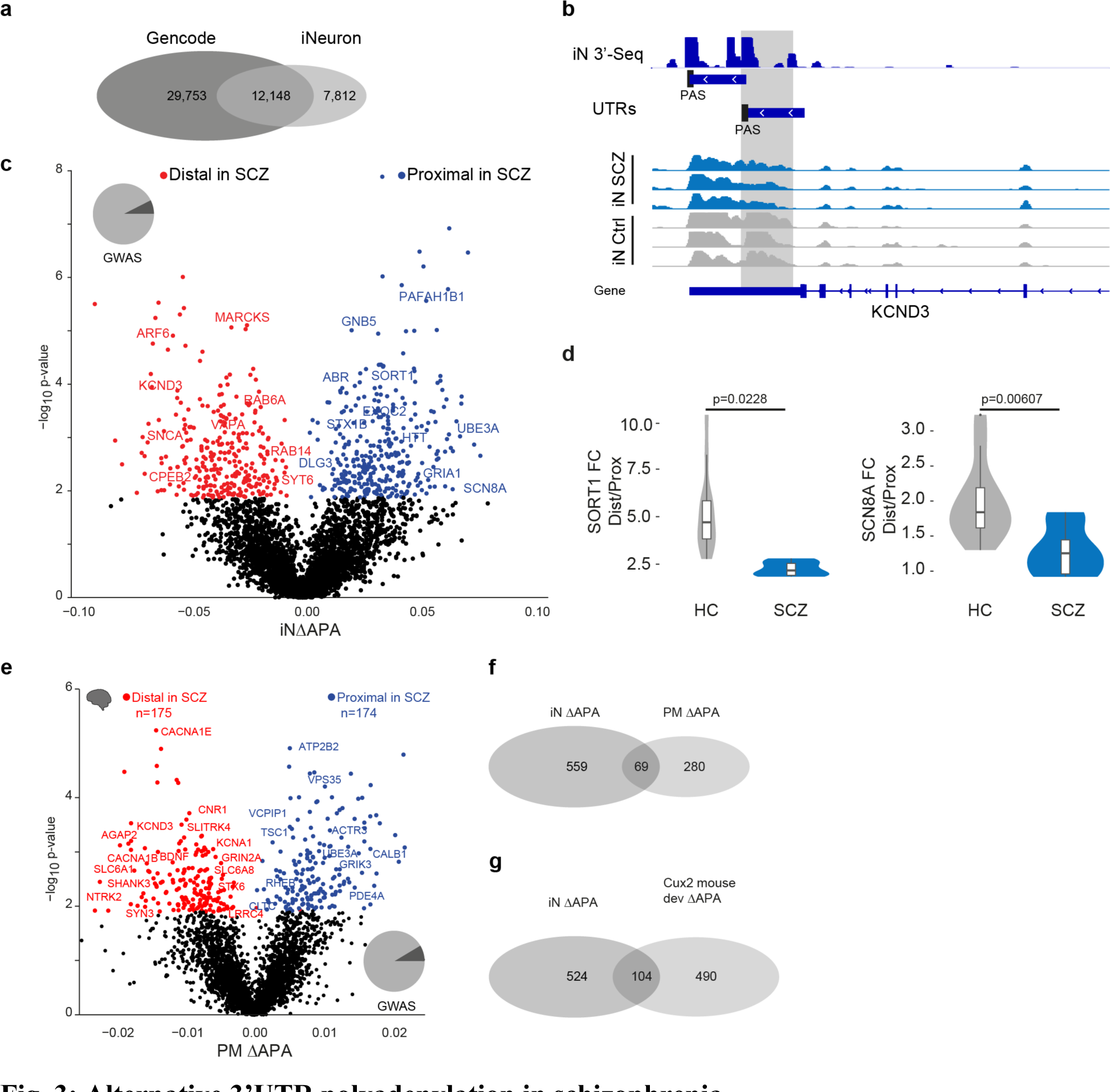
Alternative 3’UTR polyadenylation in schizophrenia. **a**, Overlap of known polyadenylation sites based on the Gencode annotation of the human genome and new polyadenylation sites identified in iNs based on 3’RNA-Seq. **b**, Example locus representation of alternative polyadenylation site detection in iNs based on 3’RNA-Seq at the KCND3 locus. From top to bottom: Read count histogram of 3’RNA-Seq in iNs cumulated across 11 donors, detected distinct UTR regions and detected proximal and distal polyadenylation sties (PAS), Read count histograms across individuals iN samples from ISCZ and HC. **c**, Volcano plot showing the difference (ΛAPA, x-axis) and significance (y-axis) of differential in 3’APA between ISCZ and HC derived iNs at day 49. Red dots indicate transcripts with a significant (FDR:<0.05) increase in usage of the distal UTR region in ISCZ, whereas blue dots delineate transcripts with a significantly higher usage of the distal UTR in the HC (higher usage of the proximal UTR in SCZ). Text highlights selected genes with dAPA **d**, Relative expression values based on qPCR for selected transcripts subjects to dAPA in n=6 iNs from ISCZ and n=6 HC samples in 2 replicates. P-values indicate significance based on LMM t-test. **e**, Volcano plot showing the difference (ΛAPA, x-axis) and significance (y-axis) of differential in 3’APA between ISCZ (n=275) and HC (n=291) in adult human postmortem PFC based on bulk RNA-Seq. Red dots indicate transcripts with a significant (FDR:<0.1) increase in usage of the distal UTR region in ISCZ, whereas blue dots delineate transcripts with a significantly higher usage of the distal UTR in the HC. **f**, Overlap of transcripts subject to dAPA between ISCZ and HC in iNs and adult PFC. **g**, Overlap of transcripts subject to dAPA between ISCZ and HC in iNs and over the course of mouse callosal projection neuron (Cux2 positive) development and maturation.

This analysis revealed widespread differences in APA, identifying 628 differential APA (dAPA) events (**Fig. 3c** and **Supplementary Table 3**) that resulted from an increased usage of either the distal or the proximal PAS to similar degree (14.29% of all genes with more than 1 polyadenylation site). In agreement with the sequencing data, qPCR confirmed dAPA for *SORT1* and *SCN8A* (**Fig. 3d**), two genes with a critical role in synapse function. In contrast, we detected virtually none dAPA events between HC and BD/MDD iNs (**Extended Data Fig. 4a**).

In order to probe the relevance of differential APA under *in vivo* conditions, we repeated the same analysis using previously published bulk RNA-Seq samples from human DLPFC in 291 controls and 275 ISCZ^11^. Consistent with the observations in iNs, this analysis confirmed widespread dAPA in postmortem PFC from ISCZ (**Fig. 3e** and **Supplementary Table 3**) as well as substantial overlap with dAPA in iNs (**Fig. 3f**). Interestingly, dAPA in iNs from ISCZ showed considerable overlap with transcripts subject to dAPA over the course of cortical development and maturation in the mouse (**Fig. 3g** and **Extended Data Fig. 4b**), suggesting a critical role of APA as one mechanism contributing to the neurodevelopmental origins of SCZ.

Strikingly, genes subject to dAPA in ISCZ were strongly enriched for synapse related processes such as glutamatergic neurotransmission, synaptic membrane components, synaptic plasticity or vesicle biology in both iNs and human PFC (**Fig. 4a**,**b**, and **Supplementary Table 3**). In particular, dAPA affected key synaptic genes such as the glutamate receptor subunits *GRIA1*, *SNCA*, *DLG3* as well as *CPLX2* critical for synaptic vesicle fusion in the forebrain^41^, the sodium channel *SCN8A* or *SORT1* involved in subcellular trafficking^42^ (**Fig. 3a** and **3e**).

**Fig. 4.**
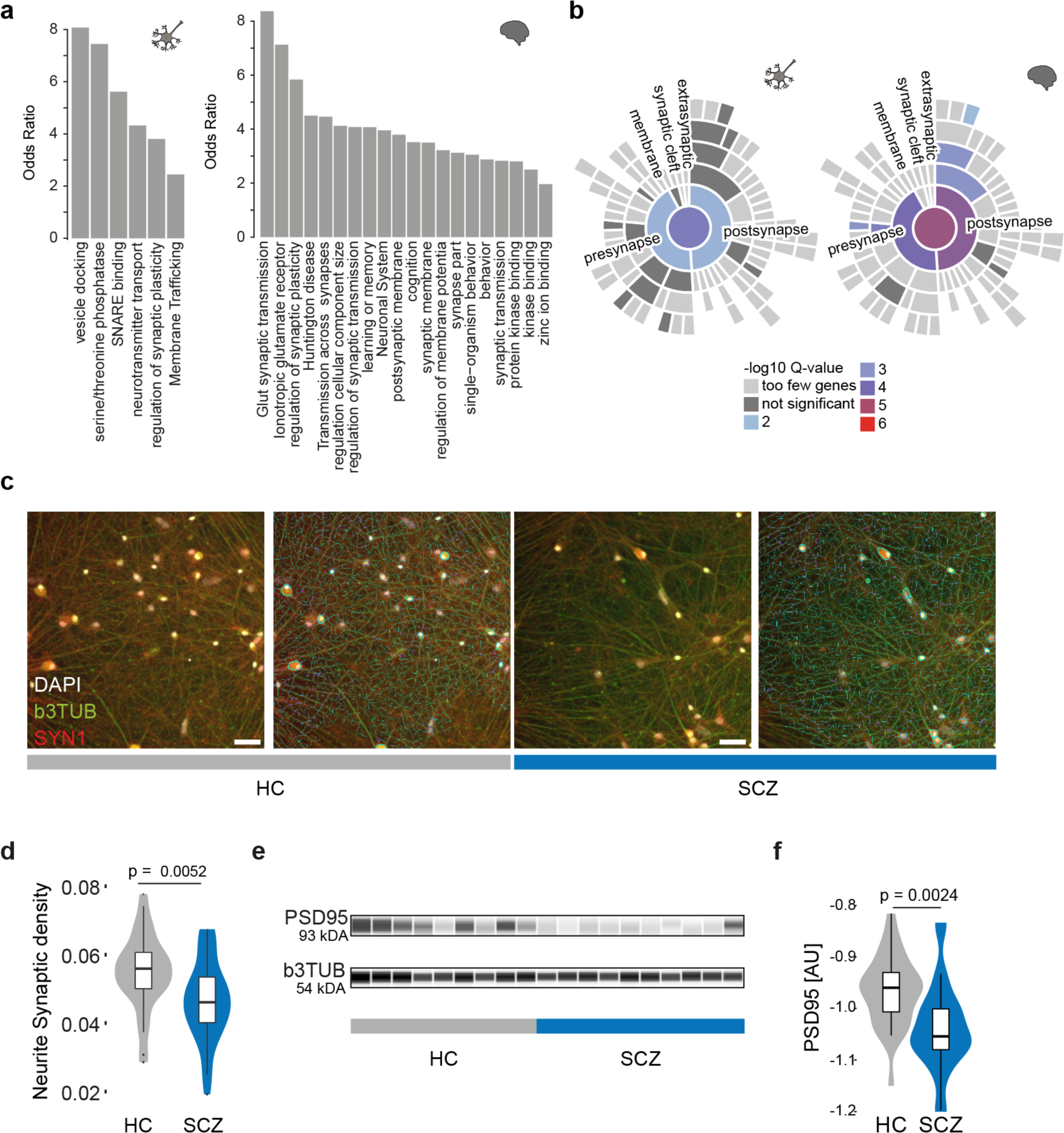
Alternative polyadenylation in schizophrenia affects synaptic pathways. **a**, Pathway enrichment (odds-ratio y-axis) for transcripts subject to dAPA between ISCZ and HC in iNs (left) or adult postmortem PFC (right). **b**, Enrichment of for transcripts subject to dAPA (between ISCZ and HC) in iNs (left) or adult PFC (right) in synapse related biological processes based on SYNGO.^46^ **c**, Representative images from high content imaging (HCI) analysis of SYN1 (red), b3TUBB (green), and DAPI (white) ICC signal without (left) and with (right) neurite segmentation mask shown in iNs from HC and ISCZ. Scale bar indicates 50 µm. **d**, Distribution of normalized SYN1 punctae density overlapping with neurites in iNs derived from HC and ISCZ donors at day 49. Density was assessed by HCI across n=10 HC and n=10 ISCZ iN samples across 66 wells in total and two independent differentiation batches. P-value indicates two-tailed significance level of linear mixed model based comparison between ISCZ and HC punctae density distributions. **e**, Representative visualization of digital quantitative western blot for PSD-95 protein abundance and b3TUB across 10 HC and 10 ISCZ iN samples at day 49 with b3TUB loading control. **f**, Distribution of normalized PSD-95 protein abundance levels in HC and ISCZ iNs at day 49 of differentiation using quantitative western blot based on the log_2_ ratio of PSD95 and bTUBB signal intensity. P-value indicates two-tailed significance level of linear mixed model based comparison of n=28 (11 donors) HC and n=26 (14 donors) SCZ samples from three independent iN differentiation batches.

To assess the functional relevance of these SCZ specific changes in APA for synapse biology, we measured synapse density by high content imaging across 20 donors (**Fig. 4c**). This analysis identified a significant reduction in synapse density (**Fig. 4d**; p-value=0.0052, LMM) between iNs derived from HC (N=10 donors) and ISCZ (N=10 donors). These results were further confirmed by quantitative western blot, revealing a discrete but significant reduction PSD95 abundance in iNs from ISCZ (**Fig. 4e,f**; p=0.0024, LMM). Jointly, these observations implicate APA as a novel mechanism contributing to alterations in synapse biology in SCZ. Importantly, consistent changes in APA across iNs and PFC suggests genetic origins of differential APA in SCZ, and support that iPSC-based models faithfully capture well-known (e.g. eQTLs) as well as until now unknown (e.g. dAPA) functional consequences of genetic factors.

## Polygenic control of differential 3’UTR APA in SCZ

To probe the link between dAPA and genetic risk in more detail, we evaluated whether or not the gene loci subject to dAPA in SCZ were enriched for SCZ associated common risk variants using partition heritability analysis^43^. This analysis revealed a significant and specific enrichment of genes affected by dAPA in iNs as well as in PFC for SCZ risk, implying a close link of APA to the genetic risk for SCZ (**Fig. 5a**).

**Fig. 5.**
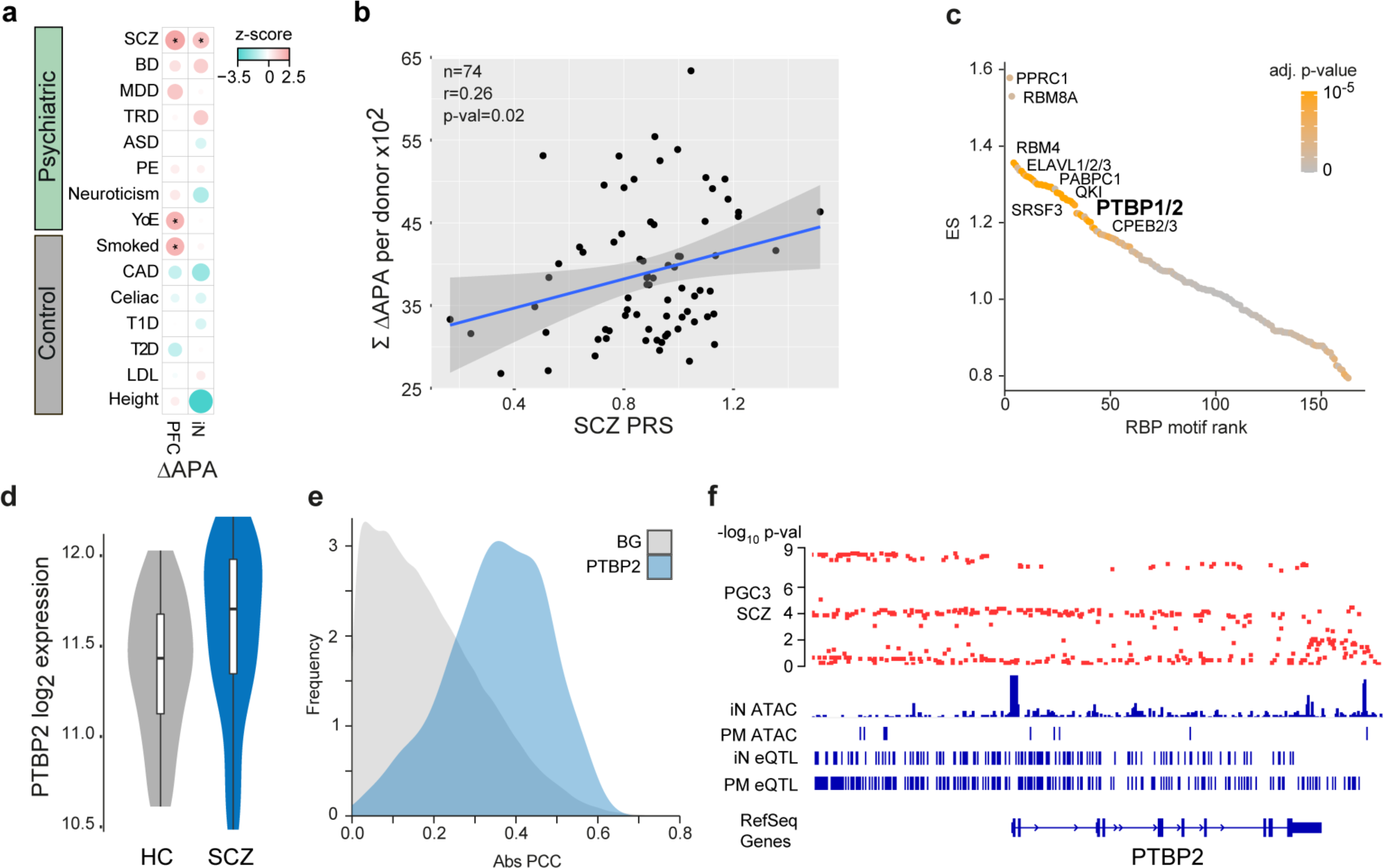
Association of dAPA with cis- and trans-acting SCZ genetic risk factors. **a**, Enrichment of trait heritability for genomic loci of genes subject to dAPA between HC and ISCZ in adult PFC and iNs for a collection of psychiatric diseases and control traits based on GWAS summary statistics and partition heritability analysis. Diseases/traits as indicated (SCZ = schizophrenia, BD = bipolar disorder, MDD = major depressive disorder, TRD = treatment resistant depression, ASD = autism spectrum disorder, PE = any psychotic experience, YoE = years of education, CAD = coronary artery disease, T1D/T2D = Type 1/2 diabetes, LDL = Levels of low-density lipoprotein). Dot size indicates enrichment levels as z-scores and * indicates significance (p-value:<0.05). **b**, Relationship of SCZ PRS (x-axis) and cumulated APA (between HC and ISCZ) in iNs across all transcripts showing differential APA (n=628) for each donor (n=74, black dots). P-value indicates significance of linear model fit (blue line) and r the corresponding pearson correlation coefficient. Grey shading signifies standard error of the fit. **c**, Distribution of RBP binding motif enrichment enrichment score (ES, y-axis) in the 3’UTR of transcripts subject to dAPA between HC and ISCZ in iNs with increased usage of the distal PAS in SCZ ordered by RBP motif enrichment rank (x-axis). Color code indicates significance of enrichment. Labeling indicates a subset of significantly enriched RBP binding motifs after multiple testing correction (adjusted p-value :<0.1). **d**, Distribution of log2 transformed variance stabilized expression level of PTBP2 in iNs from HC and ISCZ at day 49. P-value/FDR indicate significance of differential expression based on Wald-test. **e**, Frequency (y-axis) of absolute pearson correlation coefficient (PCC, x-axis) between APA level of each transcript subject to dAPA (between HC and ISCZ) in iNs (n=628) and respective PTBP2 expression level across all iN samples (blue). For comparison, results from the same analysis for all other expressed RBPs are shown as background (BG, grey). **f**, Functional genomic characterization of the PTBP2 locus in the human genome. From top to bottom: Distribution of SCZ association -log_10_ p-values based on the PGC wave 3 GWAS for all included common variants, ATAC-Seq read count histogram in iNs and PFC from postmortem (PM) human tissue, eQTLs in iNs and eQTLs in PFC.

Based on these observations, we tested the hypothesis that overall APA is a polygenically driven mechanism in SCZ, resulting from the joint action of distinct *cis*- and *trans*-acting variants. Therefore, we determined the association of cumulated APA in individual donor iN across all transcripts subject to dAPA between HC and SCZ with the respective individual polygenic risk for SCZ, BD or MDD. This analysis revealed a significant association between SCZ PRS and cumulative APA (**Fig. 5b**; r=0.26, p=0.02) but not BD or MDD PRS (**Extended Data Fig. 4c,d**), suggesting polygenic origins of dAPA consistent with overall GWAS signal enrichment (**Fig. 5a**). In contrast, association of cumulated APA across transcripts subject to dAPA between HC and BD/MDD using relaxed statistical inclusion criteria (q-value:<0.1) did not show any association with any PRS (**Extended Data Fig. 4e-g**).

To pinpoint specific genetic variants contributing to dAPA, we evaluated *cis*-acting common SNPs for their potential to operate as APA qTLs. This analysis revealed no associations, likely due to lack of power. Given the importance of *trans*-acting factors as potential amplifiers of *cis*-regulatory variants and the previously reported role of RNA binding proteins (RBPs) in the genetic basis of mental disorders^17^, we sought to identify de-regulated RBPs as potential causes of dAPA in iNs from ISCZ. RBP motif analysis in the UTRs of transcripts subject to dAPA identified numerous expressed RBPs with critical roles for neuronal biology such as QKI, ELAVL and RBM proteins (**Fig. 5c** and **Supplementary Table 3**). However, only PTBP2 and PCBP2 were also differentially expressed between iNs from HC and ISCZ. PTBP2 had previously been implicated in APA^44^ and differential expression in iNs (p=3.24×10^-5^; **Fig. 5d**) extended to postmortem PFC (p=0.018) when comparing HC and ISCZ. Importantly, PTBP2 expression showed a high correlation to APA of transcripts subject to dAPA in iNs (**Fig. 5e**, blue) compared to APA of randomly selected transcripts (**Fig. 5e**, grey, and **Extended Data Fig. 5**).

In addition, PTBP2 is located in a SCZ GWAS risk locus (**Extended Data Fig. 5a**) which harbors multiple regulatory elements in iNs and PFC and exhibits a significant eQTL in both iNs and PFC (**Fig. 5f**), indicating de-regulation of PTBP2 by *cis*-acting SCZ associated genetic variation (**Extended Data Fig. 5b**). However, in light of the low difference in risk allele frequency distribution of the PTBP2 eQTL, *cis*-acting genetic effects can only explain a fraction of PTBP2 de-regulation in SCZ (R^2^_rs72721996_ =0.065). However, in addition to cis-acting genetic variation, PTBP2 mRNA was predicted to be targeted by the SCZ associated microRNA let-7-5p^45^ (**Fig. 2b**) acting in trans. In line with this prediction, we observe an anti-correlation of PTBP2 expression with miR-let-7-5p (R_Pearson_=-0.318) and a significant increase in explained variance of overall PTBP2 expression (R^2^_rs72721996+let-7-5p_= 0.1901) based on a linear model incorporating donor genotype and let-7-5p expression (p=0.00257, **Extended Data Fig. 5c**). Jointly, these observations support the notion that multifactorial de-regulation of PTBP2 expression levels partially mediates dAPA in neuronal cells from ISCZ.

Based on these findings, we evaluated whether or not PTBP2 contributes to dAPA in iNs from ISCZ through shRNA mediated acute knockdown (**Fig. 6a**,**b**). RNA-Seq based assessment of dAPA between PTBP2*^KD^* and scramble control in iNs revealed widespread dAPA (**Fig. 6c**; n=546), biased towards usage of the more proximal polyadenylation site in the KD condition (n=432 vs n=114). Affected transcripts showed substantial overlap with dAPA between HC and ISCZ in iNs as well as PFC (**Fig. 6d**), suggesting a contribution of PTBP2 to dAPA in these conditions. Similar to the conditions in iNs and PFC from ISCZ, transcripts affected by dAPA upon PTBP2 knockdown were enriched for various synapse related processes (**Fig. 6e**).

**Fig. 6.**
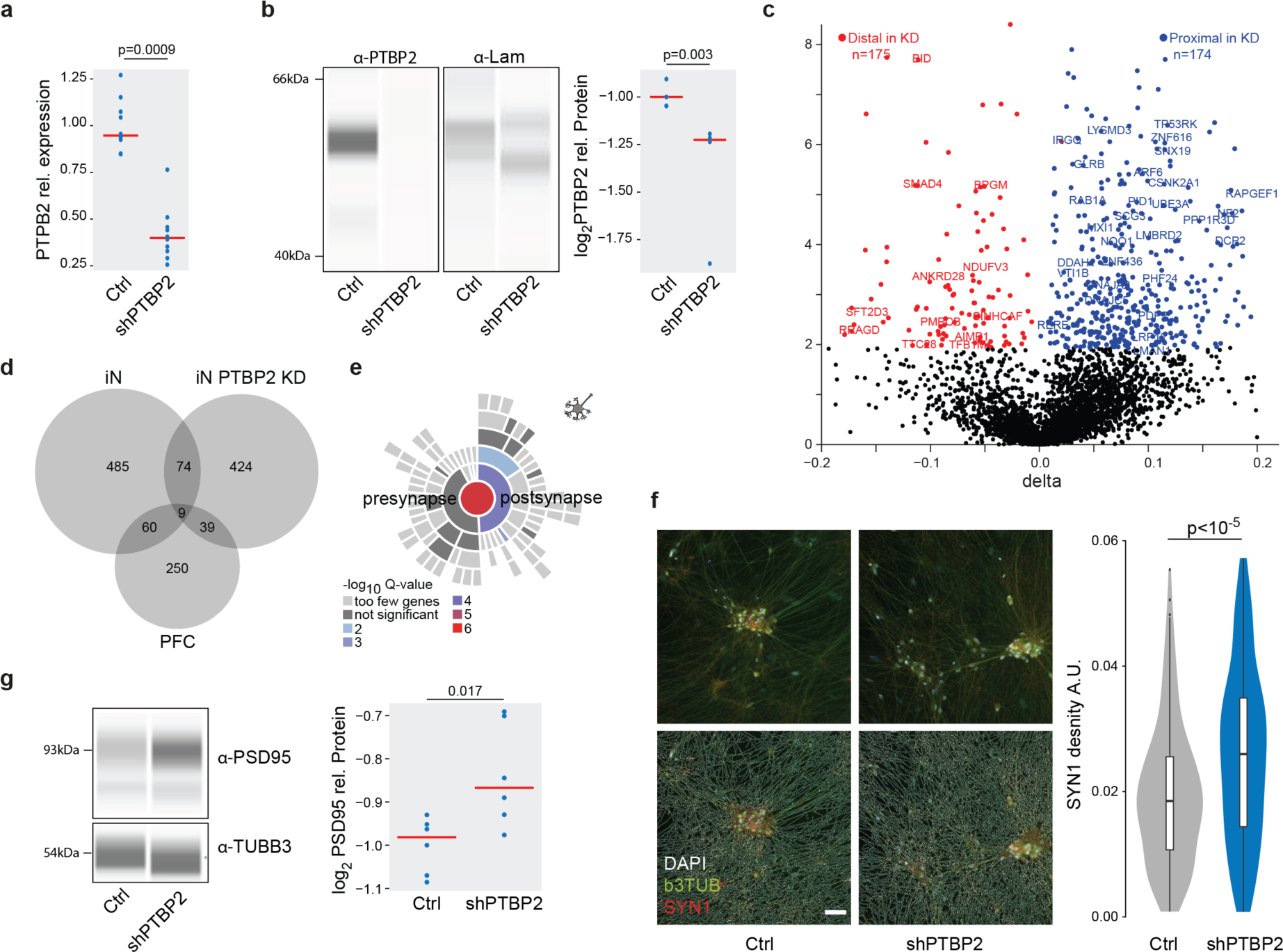
PTBP2 is a trans-acting genetic risk factor for SCZ mediating dAPA dependent synaptic density alterations. **a**, Distribution of relative expression levels (y-axis) of PTBP2 normalized to the housekeeping gene RTF2 in HC derived iNs treated with scramble control and a pool of 4 PTBP2 shRNAs based on qPCR. P-value indicates two-tailed significance level of linear mixed model based comparison of n=20 samples from 3 distinct donors and three independent iN differentiation batches correcting for batch as random effect. **b**, Left: Representative visualization of digital quantitative western blot for PTBP2 protein abundance and laminin in scramble control and PTBP2 shRNA mediated knockdown in iNs. Right: Distribution of normalized PTBP2 protein abundance levels (y-axis) in scramble control and PTBP2 shRNA mediated knockdown in iNs based on quantitative western blot. P-value indicates two-tailed significance level of linear mixed model based comparison of n=15 samples from 3 distinct donors and three independent iN differentiation batches correcting for batch as random effect. **c**, Volcano plot showing the difference (ΔAPA, x-axis) and significance (y-axis) of dAPA between scramble control (n=2) and acute shRNA mediated knockdown of PTBP2 at the end of the differentiation for 5 days (n=2) in iNs. Red dots indicate transcripts with a significant (FDR:<0.05) increase in usage of the distal UTR region in the knockdown condition, whereas blue dots delineate transcripts with a significantly higher usage of the distal UTR in the scramble control. **d**, Overlap of transcripts subject to dAPA between HC and ISCZ in iN or PFC or between scramble control and PTBP2 knockdown in iNs. **e**, Enrichment for transcripts subject to dAPA between scramble control and PTBP2 knockdown in iNs in synapse related biological processes based on SYNGO^46^. **f**, Left: Representative image of SYN1 (red) distribution in iNs infected with scramble control shRNA or PTBP2 knockdown alongside with bTub (green) and DAPI (white) signal without (upper line) and with (top line) neurite segmentation mask. Right: Distribution of SYN1 punctae area density (arbitrary units) in neurites of iNs with scramble control or PTBP2 knockdown in iNs from 3 different donors each, 23 wells and 291 fields of view. P-value indicates two-tailed significance level of linear mixed model based testing correcting for donor as random effect. Scale bar indicates 50 µm. **g**, Left: Representative visualization of digital quantitative western blot for PSD-95 protein abundance and b3TUB in scramble control and PTBP2 shRNA mediated knockdown in iNs. Right: Distribution of normalized PSD-95 protein abundance levels for high molecular weight (95 kDa) (y-axis) in scramble control and PTBP2 shRNA mediated knockdown in iNs based on quantitative western blot. P-value indicates two-tailed significance level of linear mixed model based comparison of n=10 samples from 3 distinct donors and three independent iN differentiation batches correcting for batch as random effect.

Subsequent assessment of synaptic density in PTBP2*^KD^*iNs revealed a significant increase in synapse density (**Fig. 6f**) and PSD-95 protein level (**Fig. 6g**). These results corroborate a decrease in synaptic density and PSD-95 levels upon higher expression of PTBP2 in iNs and PFC from ISCZ, supporting a causal role of PTBP2 in mediating APA dependent changes in synaptic components.

Jointly, these findings establish dAPA as molecular mechanism through which diverse SCZ associated genetic variants contribute to synaptic changes in disease relevant glutamatergic neurons of the brain. Moreover, our results demonstrate convergence of diverse *cis*- and *trans*-acting genetic variants specific to SCZ on one common molecular mechanism.

## Discussion

In this study, we tested the hypothesis whether polygenic and heterogenous genetic risk for psychiatric disease does functionally converge across patients onto common molecular mechanisms that regulate genes with a central role in disease relevant processes.

To that end, we employed the largest iPSC cohort in the field of psychiatry until today to unravel the functional consequences of distinct polygenic risk factor combinations. This endeavor identified numerous new as well as previously implicated genes and microRNAs associated with SCZ or BD in iPSC-derived neurons.

However, the majority of the identified genes are not located in loci identified by SCZ GWAS and do not seem to be modulated by single *cis*-acting regulators. Instead, the expression of these genes is rather governed by a complex matrix of multiple *cis*- and *trans*-acting factors. Given the minimal differences in disease associated allele frequency of GWAS hits and corresponding heterogeneity in genetic risk factor combinations across the patient population, this is not surprising. Against this background, it is highly unlikely to observe consistently altered genes in GWAS loci, if their effect is expected to be caused predominantly by *cis* acting CVs.

In this study, diagnosis associated changes in gene expression, microRNA expression and open chromatin in iNs showed limited enrichment for specific biological processes or genetic risk. Instead, we identify APA as new mechanism contributing to the molecular basis of SCZ, particularly affecting synaptic genes. Importantly, APA in iNs and PM is significantly associated with SCZ polygenic risk, suggesting a substantial *cis*-acting contribution of genetic risk from common variants to dAPA. Moreover, the consistent alteration of many transcripts across donors with diverse polygenic risk factor combinations implies the presence of functional convergence of these risk factors through dAPA. One potential mechanism that can mediate such functional convergence is the de-regulation of a *trans*-acting regulator of APA such as PTBP2. In this context, the alteration of PTBP2 expression can be driven by various causes such as *cis*-acting SCZ associated genetic variation or other *trans*-acting mechanisms (e.g. microRNAs) depending on the complex genetic background of each donor. The functional consequences of its de-regulation however result in the consistent APA of many synaptic transcripts across ISCZ, partially explaining the observed variation in APA in iNs and PFC.

We show that PTBP2 de-regulation directly impacts synapse density in iNs, with the directional effect consistent with the observation in ISCZ as one functional consequence of APA. It is well established that APA plays a key role in modulating mRNA half lifetime, translation efficiency as well as subcellular localization. Thus, given the large number and diversity of transcripts affected by APA, we hypothesize that APA in ISCZ leads to alterations in distinct physiological properties of neuronal cells, both in the adult brain and over the course of development.

Overall, our findings suggest that distinct genetic risk factor combinations in ISCZ control *cis*- and *trans*-acting regulators that converge on 3’UTR APA as one critical molecular mechanism that drives de-regulations of downstream effector genes with a prominent role in synaptic function. In perspective, our findings also offer conceptual insight into the long-standing question of how heterogeneous patient populations can share clinical diagnosis based on converging molecular psychopathology.

While this study focuses on SCZ, we further propose that deep phenotyping of well-powered iPSC studies can provide in general new insights into those molecular mechanisms, downstream effector genes, and biological processes on which polygenic risk converge and translates into phenotypic states. Such insights will be critical to narrow today’s gap in genotype-phenotype relationships in complex diseases and to identify new therapeutic entry points.

## Extended Data Figures

**Extended Data Fig. 1.**
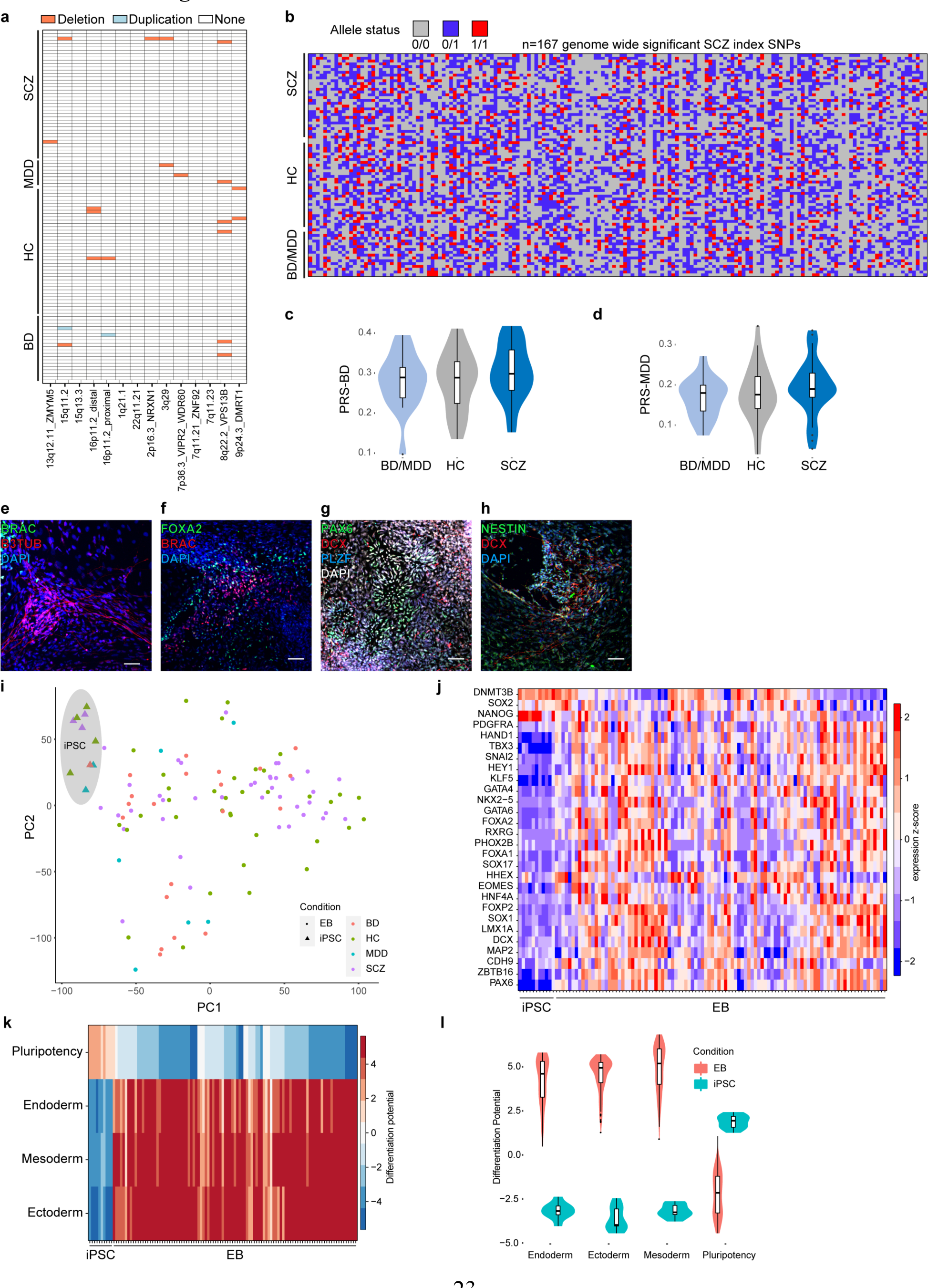
Quality control of iPSCs, and genetic characterization. **a**, Distribution of previously identified CNVs associated with SCZ (x-axis) across all iPSC lines ordered by diagnosis group (y-axis). **b**, Distribution of index-SNP allele status (color-code) for 167 genome wide significant SCZ associated loci (x-axis) across all iPSC donors. **c**, Distribution of the individual polygenic risk score for bipolar disorder for the different diagnosis groups of hiPSC donors. **d**, Distribution of the individual polygenic risk score for major depressive for the different diagnosis groups of hiPSC donors. **e**, Representative immunofluorescence image of an embryoid body section at day 15 of differentiation colored for mesodermal marker Brachury (BRAC) and ectodermal marker beta-3-tubulin (B3TUB). Scale bar indicates 50 µm. **f**, Representative immunofluorescence image of an embryoid body section at day 15 of differentiation colored for endodermal marker FOXA2 and mesodermal marker BRAC. Scale bar indicates 50µm. **g**, Representative immunofluorescence image of an embryoid body section at day 15 of differentiation colored for ectodermal markers PAX6, DCX and PLZF. Scale bar indicates 50µm. **h**, Representative immunofluorescence image of an embryoid body section at day 15 of differentiation colored for ectodermal markers NESTIN and DCX. Scale bar indicates 50µm. **i**, PCA analysis results for RNA-Seq profiles from EBs derived from all iPSC lines (stars) and a subset of iPSC analyzed in the pluripotent state (triangles) colored by diagnosis group. **j**, Heatmap of key lineage marker gene expression (y-axis) across all EBs and iPSCs analyzed in this study (y-axis) as z-scores (color scheme) based on RNA-Seq. **k**, Cumulated differentiation potential for each germ layer and pluripotency signature assessed (y-axis) by RNA-Seq for EBs (see methods for details) across iPSC lines (x-axis). **l**, Distribution of cumulated differentiation potential signature (y-axis) for each germ layer and the pluripotency signature and (x-axis) across iPSC-derived EBs (red) and a subset of iPSCs (blue).

**Extended Data Fig. 2.**
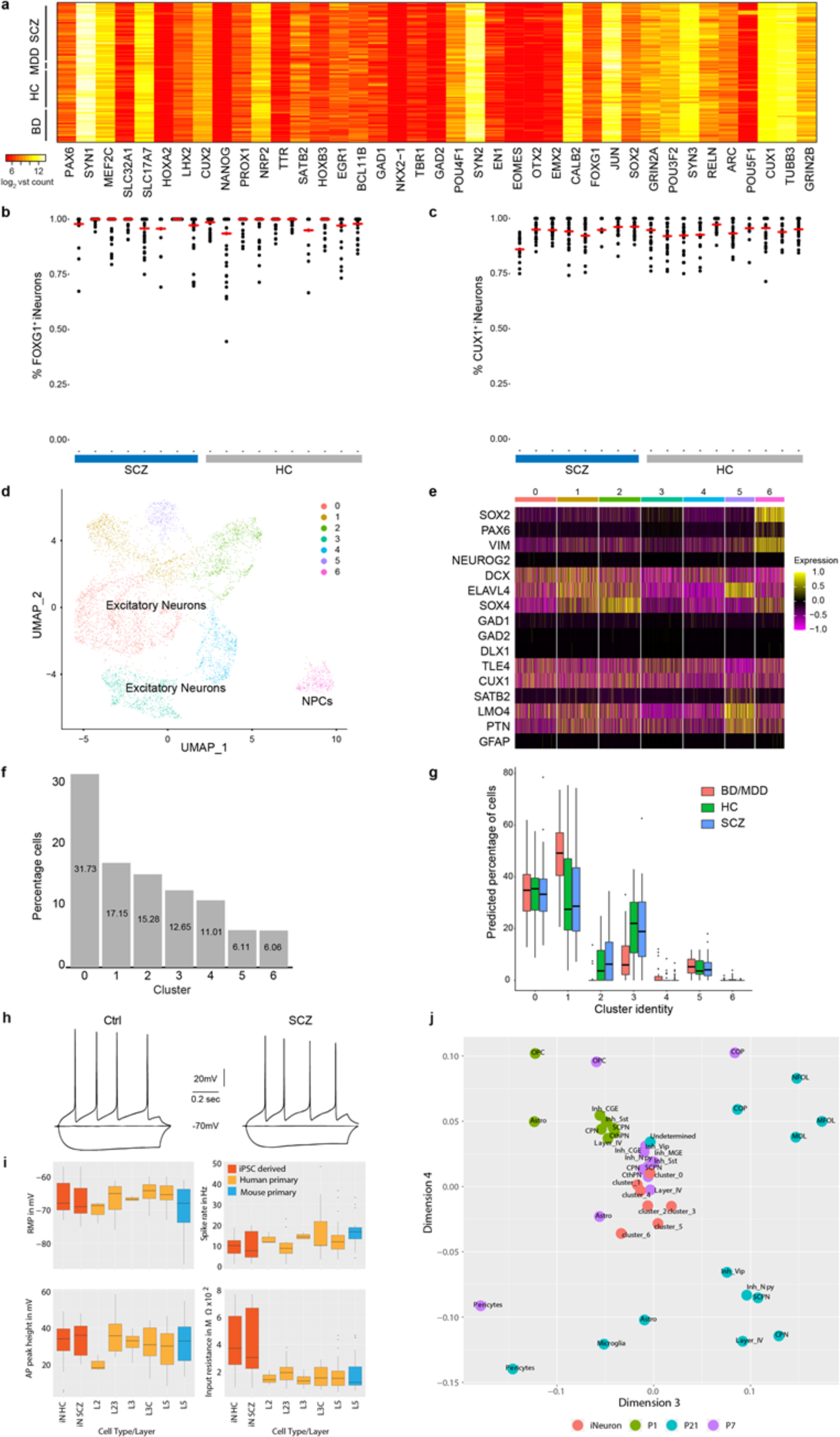
Characterization and quality control of multi-omics iPSCderived neuron profiles. **a**, Expression of key neuronal marker genes (x-axis) across all interrogated iPSC lines (y-axis) in iPSC derived neurons at day 49 based as log_2_ transformed RNA-Seq counts. **b**, Dot plot showing FOXG1+ iNs after 49 days of differentiation from HC (n=10, grey) and SCZ (n=8, blue) based on ICC analysis. Each dot represents one FOV. Red lines indicate median of FOXG1+ iNs per cell line. **c**, Dot plot showing CUX1+ iNs after 49 days of differentiation from HC (n=10, grey) and SCZ (n=8, blue) based on ICC analysis. Each dot represents one FOV. Red lines indicate median of CUX1+ iNs per cell line. **d**, Umap representation of iNs single-cell RNA-Seq profiling. Cell type identities were assigned based marker gene expression. Cluster 6 represent neural progenitor cells (NPCs). **e**, Heatmap for subsampled 1000 cells from a. showing the SCT scaled expression of selected marker genes across iN clusters (x-axis). Note, the increased expression of NPC markers(SOX2, PAX6, VIM) in cluster 6 and the absence of interneuronal marker gene expression (GAD1/2, DLX1) and GFAP expression in all clusters. **f**, Percentage of cells (y-axis) per cluster (x-axis) from single-cell RNA-Seq of iNs from a. **g**, Distribution of predicted cell identity proportion (y-axis) per single cell RNA-Seq cluster (x-axis) for each iN bulk RNA-Seq dataset splitted by diagnoses (colored boxplots). Predicted proportions were determined through in silico deconvolution analysis using the scRNA-Seq dataset from iNs as reference. Boxes indicate 25^th^ and 75^th^ quantiles, black bar median, lines demarcate 1.5 IQR from the median in either direction. **h**, Representative traces for HC and SCZ iNs of the electrophysiological basic characterization by patch-clamp recordings. **i**, Comparison of basic electrophysiological properties from single cell patch-clamp recordings of iNs from HC or ISCZ (red) to previously published data^28^ from primary human (orange) or mouse neurons (blue). X-axis indicates layer of origin of patched cells. RMP – resting membrane potential in mV, AP peak height – peak height of action potential relative to baseline (overshoot) in mV, Spike rate – frequency of action potentials in Hz calculated from 700/800ms measurement windows, Input resistance – whole cell input resistance in mega Ω. L2/3/5 indicates layer of neuron origin. **j**, Multidimensional scaling analysis comparing pseudo-bulk profiles of scRNA-Seq from the developing mouse cortex at developmental stages P1, P7 and P21 and individual transcriptomic profiles of all clusters detected in the iN dataset. OPC – oligodendrocyte precursor; COP, NFOL, MFOL, MOL – distinct stages of oligodendrocytes; CPN, ScPN, CThPN - excitatory neuronal projection neuron subtypes, Inh-interneuron subtypes. Cluster indicate iNeuron clusters from a.

**Extended Data Fig. 3.**
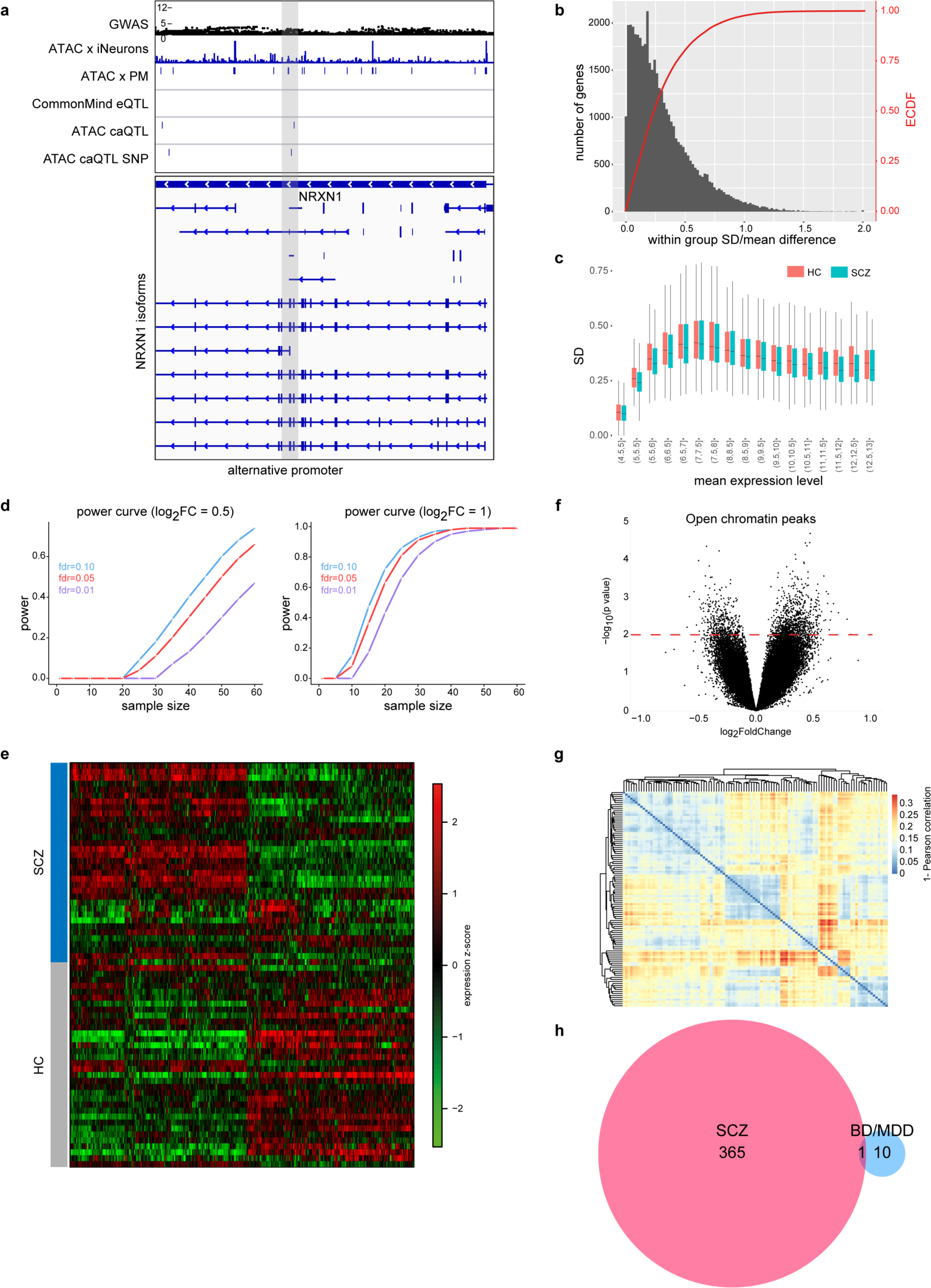
Differential expression analysis and power calculations of iPSC-derived neurons. **a**, Representative example of genomic annotations and distinct qTL types at the intronic region of NRXN1 locus, from top to bottom: Association strength of imputed SNPs at the locus with SCZ based on SCZ GWAS, ATAC-Seq profile of representative iPSC derived neuron at day 49, eQTLs detected at the locus in iNs, caQTLs detected at the locus co-localizing with open chromatin regions. Grey vertical bar indicated an alternative promoter region of NRXN1 in which a caQTL is located. **b**, Frequency of genes (y-axis left) characterized by a particular ratio of within group standard deviation (SD) over mean group difference (HC vs SCZ, x-axis) as well as empirical cumulative density function (ECDF) of genes (red-curve, y-axis right). **c**, Distribution of standard deviation (y-axis) for all genes within a certain mean expression level interval (x-axis) per sample group (color code). Boxes indicate 25^th^ and 75^th^ quantiles, black bar median, lines demarcate 1.5 IQR from the median in either direction.**d**, Power estimation (y-axis) to detect differentially expressed genes between two sample groups with individual sample size per group indicated on the x-axis at distinct FDR thresholds indicated by colored lines. Power estimates were conducted using a negative binomial model with empirically inferred dispersion parameters using the iPSC iN RNA-Seq profiles and a minimum effect size of log_2_ fold-change 0.5 (left) or 1 (right). **e**, Heatmap showing differentially expressed genes (DEG) (FDR≤0.01 and |log_2_FC|≥0.4 x-axis) between HC and ISCZ iNs (y-axis) as z-scores (color code). **f**, Volcano plot showing the log_2_ fold change (x-axis) and significance (p-value) of differential open chromatin signal between SCZ and HC iNs at day 49. No peaks pass the significance threshold (FDR:<0.05). Dashed red line indicates nominal significance. **g**, Clustered heatmap of ATAC-Seq peak profiles of all iNs from this study using 1-pearson correlation coefficient as metric (color code). **h**, Overlap of DEGs between SCZ vs HC iNs (red) and BD/MDD vs HC iNs (blue).

**Extended Data Fig. 4.**
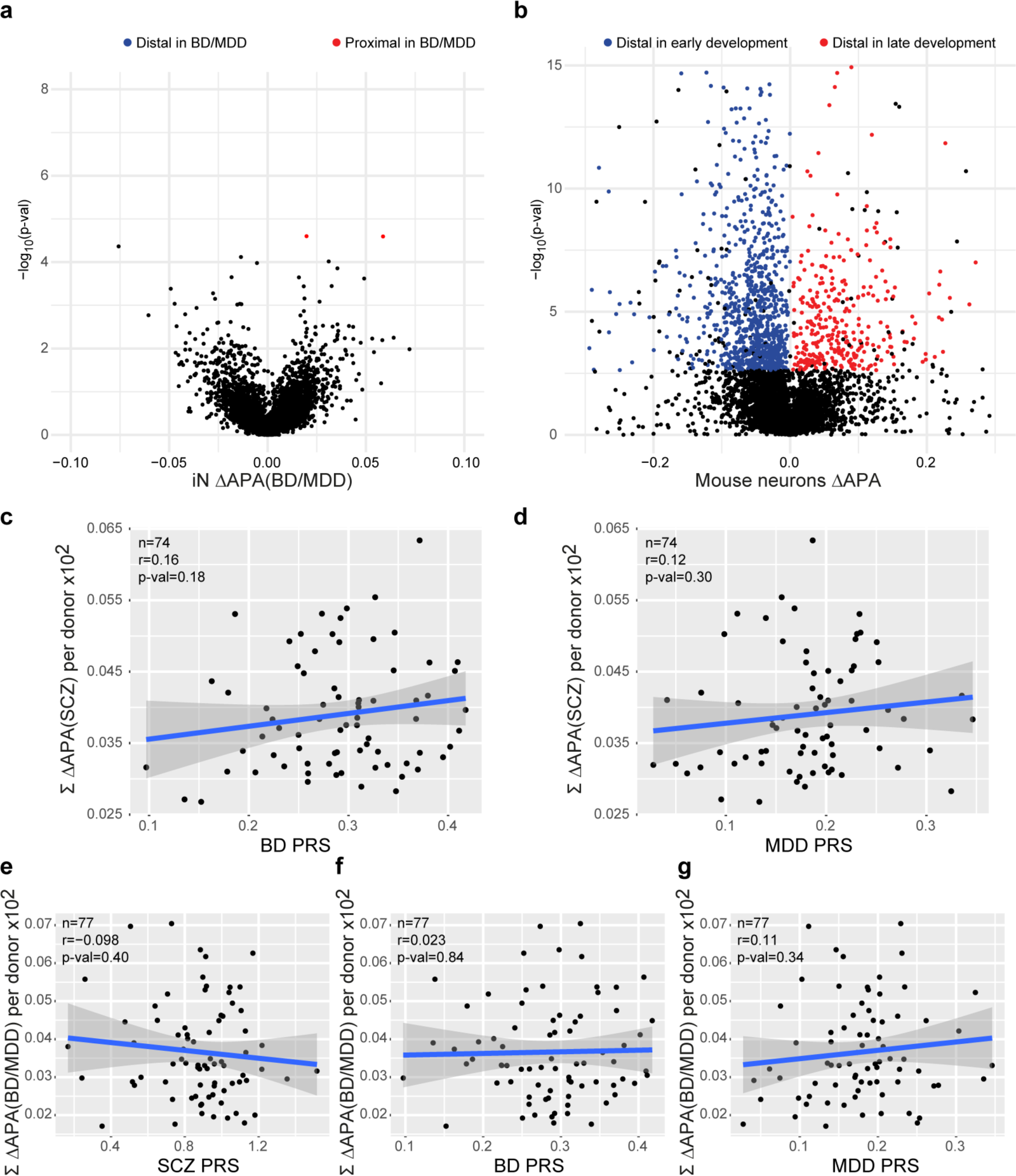
dAPA in bipolar disorder, mouse neurons, and its associations with genetic risk. **a**, Volcano plot showing the difference (ΔAPA, x-axis) and significance (y-axis) of differential in 3’APA between BD/MDD and HC derived iNs at day 49. Red dots indicate transcripts with a significant (FDR:<0.05) increase in usage of the distal UTR region in HC (higher usage of the proximal UTR in SCZ), whereas blue dots delineate transcripts with a significantly higher usage of the distal UTR in the BD/MDD. **b**, Volcano plot showing the difference (ΔAPA, x-axis) and significance (y-axis) of dAPA between mouse callosal projection neurons during early development (E18.5-P1) and late developmental stages (P21-P48). Red dots indicate transcripts with a significant (FDR:<0.05) increase in usage of the distal UTR region in at late developmental stages, whereas blue dots delineate transcripts with a significantly higher usage of the distal UTR during the early phases of postnatal development **c**, Relationship of BD PRS (x-axis) and cumulated APA (between HC and ISCZ) in iNs across all transcripts showing differential APA (n=628) for each donor (n=74, black dots). P-value indicates significance of linear model fit (blue line) and r the corresponding pearson correlation coefficient. Grey shading signifies standard error of the fit. **d**, Relationship of MDD PRS (x-axis) and cumulated APA (between HC and ISCZ) in iNs across all transcripts showing differential APA (n=628) for each donor (n=74, black dots). P-value indicates significance of linear model fit (blue line) and r the corresponding pearson correlation coefficient. Grey shading signifies standard error of the fit. **e**, Relationship of SCZ PRS (x-axis) and cumulated APA (between HC and BD/MDD) in iNs across all transcripts showing differential APA (n=20) for each donor (n=77, black dots). P-value indicates significance of linear model fit (blue line) and r the corresponding pearson correlation coefficient. Grey shading signifies standard error of the fit. **f**, Relationship of BD PRS (x-axis) and cumulated APA (between HC and BD/MDD) in iNs across all transcripts showing differential APA (n=20) for each donor (n=77, black dots). P-value indicates significance of linear model fit (blue line) and r the corresponding pearson correlation coefficient. Grey shading signifies standard error of the fit. **g**, Relationship of MDD PRS (x-axis) and cumulated APA (between HC and BD/MDD) in iNs across all transcripts showing differential APA (n=20) for each donor (n=77, black dots). P-value indicates significance of linear model fit (blue line) and r the corresponding pearson correlation coefficient. Grey shading signifies standard error of the fit.

**Extended Data Fig. 5.**
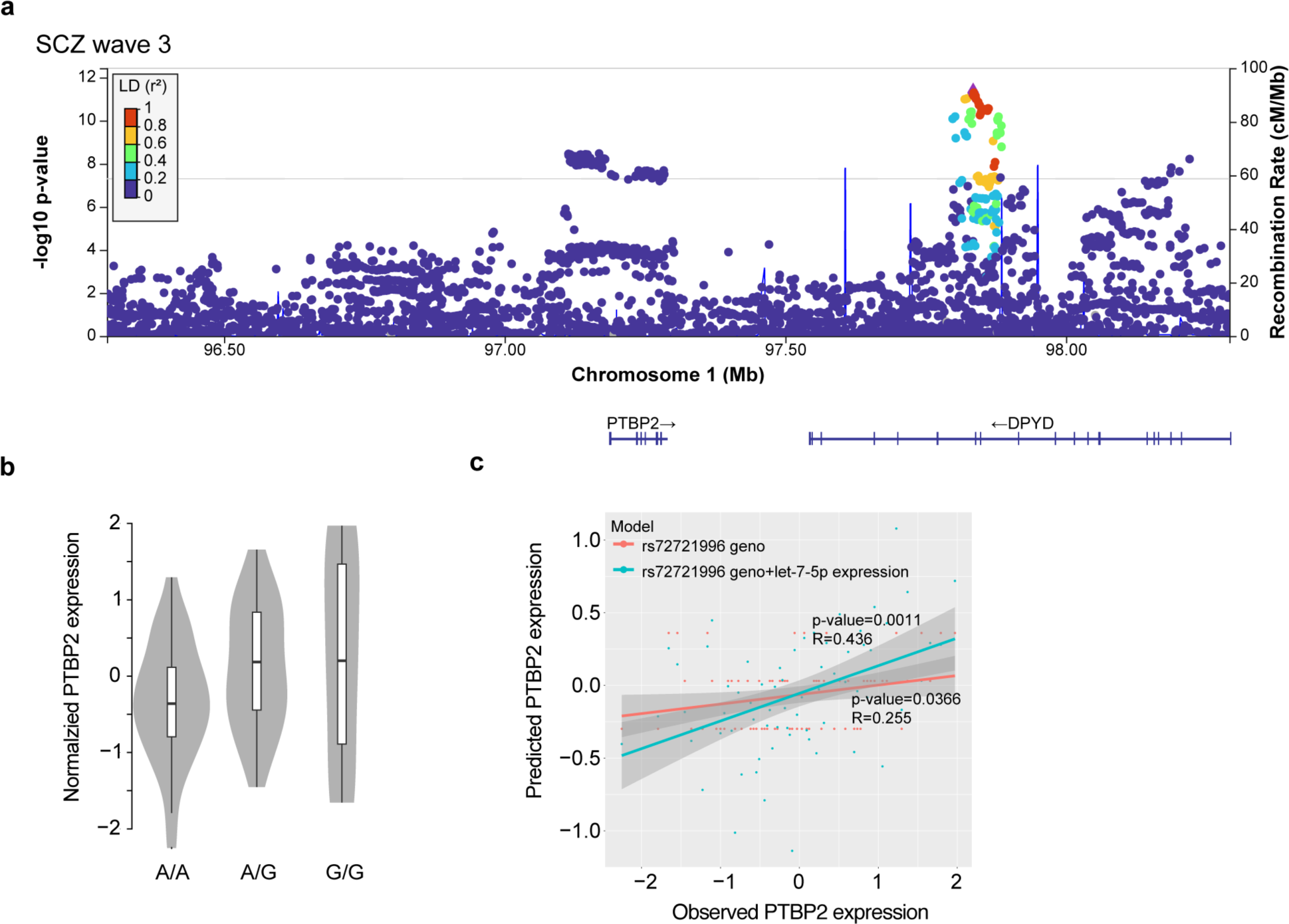
Characterization of the PTBP2 gene. **a,** Locus zoom plot of PTBP2 locus showing linkage disequilibrium (LD) scores and -log_10_ p-value based on SCZ GWAS from the PGC wave 3. **b,** Normalized PTBP2 expression levels in iNs from all donors as a function of allele status at the PTBP2 eQTL rs72721996 at the SCZ risk locus PTBP2. **c,** Results of linear model-based regression of PTBP2 expression levels in iNs on donor genotype only (red) or donor genotype + miR-let-7-5p expression in the same donors across n=67 samples. X-axis depicts measured PTBP2 expression levels while y-axis shows predicted expression values based on the respective model. P-values indicate significance of the respective model while R values show overall correlation of measured and predicted expression values.

## Supplementary Tables

**Supplementary Table 1. Overview of included cell lines**

Explanation of the columns from left to right: Classifier of diagnosis; Treatment information indicates recorded lithium treatment in BD patients and if patients with SCZ received clozapine treatment or treatment with other antipsychotic medication (noClozapin); Sex indicates male (M) and female (F) donors; age at timepoint of donation; Age at (recorded) onset of diseases; hiPSC generator; donor tissue; reprogramming method; RNAseq, ATAC-Seq and microRNA-Seq.

Separate Excel file.

**Supplementary Table 2. Summary of eQTL, caQTL and microRNA qTLs identified in this study**

Separate Excel file.

**Supplementary Table 3. Summary of differentially expressed genes (DEGs), microRNAs (DiffmicroRNA) and differential APA analysis (dAPA)**

Separate Excel file.

**Supplementary Table 4. Summary of iN endophenotyping data used for statistical analysis**

Separate Excel file.

**Supplementary Table 5. Summary of key reagents used**

Separate Excel file.

**Supplementary Data 1. Quality control of iPSC genotype**

This data summarizes the digital karyotyping results of all iPSC lines used in this study using GSA genotyping array. For each line, a pairwise CNV analysis was conducted comparing the iPSC genotype to that of the original donor derived from a donor’s blood sample. The top plot for each donor summarizes detected CNVs between iPSC and blood based genotype, where blue lines indicate a CNV. The two plots below show the CNV status in the donor sample (top) and iPSC line (bottom), where the color code indicates the type of CNV found in each sample.

Separate PDF file.

**Supplementary Data 2. Quality control of iPSC pluripotency markers**

This data figures summarizes representative immunocytochemistry images for the key pluripotency markers OCT4, TRA1-60, NANOG, SOX2 as well as differentiation related markers PAX6, SOX1, NESTIN and KI-67 across all iPSC lines used in this study, highlighting robust pluripotency marker presence and absence of differentiation related markers. Pages 1-13, scale bar indicates 100µm; pages 14-24, scale bar indicates 50µm; pages 25 and 26, scale bar indicates 100µm. Separate PDF file.

**Supplementary Data 3. Raw quantitative western blot images.**

Fully pictured digital quantitative western blot performed with WES protein simple. Western blot for each epitope was performed in 3 independent iNeuron differentiation batches from Ctrl (grey) and SCZ (blue) at day 49. Test lanes for subsequent assay optimization, e.g. concentration optimization, are not excised but marked with black bars above the respective WB lane.

Abundance of PSD95 and b3tub for normalization, is illustrated in classical WB-like lane view above the graph view with detected chemiluminescence on the y-axis and molecular weight (MW) on the x-axis, which is the base for digital quantification. Standardized ladders illustrate the molecular weight of 12, 40, 66, 116, 180, 230 kDA. Separate PDF file.

## Experimental Model and Subject details

All tissue donations from patients with schizophrenia (SCZ), bipolar disorder (BD) and depression (MDD) and neurotypical controls (Ctrls) were conducted according to the guidelines of the Declaration of Helsinki. Informed consent was obtained from all subjects involved in the study. Human iPSC cell lines were obtained from cohort studies at the Department of Psychiatry and Psychotherapy, University Hospital, LMU Munich, 80336 Munich, Germany (approved by the local ethics committee of the Faculty of Medicine, LMU Munich, Project 17-880, 29.03.2018; project 18-716, 15.10.2020) and at the MPI for Psychiatry (approved by the local ethics committee of the Faculty of Medicine, LMU Munich, project numbers 350-14, 19-310, 20-314,19-678 and 18-393 and from ThermoFisher, NIMH, Helmholtz Zentrum Munich and the Coriell Biobank. From the latter, we also included a cohort of Amish individuals suffering from BD/MDD and respective controls, see **Supplementary Table 1**.

## Key reagents and resources used in this study

**Supplementary Table 5**

## PBMC collection and iPSC reprogramming

### PBMC collection at LMU

Blood samples (18-36 ml) were collected in EDTA or natrium heparin tubes. PBMCs were isolated by Ficoll-Paque gradient according to manufacturer’s instructions. After washing with PBMC medium (RPMI1640, 10% FBS, and 1 mM sodium pyruvate), cells were counted, and resuspended (1×10^7^ cells/ml) in freezing medium (RPMI1640, 20% FBS, 1 mM sodium pyruvate, and 10% DMSO). Cryotubes were placed immediately into a freezing container, stored at -80°C for 24 hr, and thereafter under liquid nitrogen.

### Reprogramming of PBMCs to iPSCs at LMU

Reprogramming was carried out by modification of a previously published protocol^47^. PBMCs were thawed in pre-warmed StemPro34 medium (StemCell Technologies^TM^) and cultivated in uncoated 6 well plates for 5 days (D). Half media change was performed on D 1 and D 4.

After 2 washes with DPBS, PBMCs (2 x 10^6^) were resuspended in 100 µl nucleofection buffer (Human CD34+ Nucleofector Kit^TM^), episomal vectors^47^ were added (pCE-hOCT3/4 (#41813), pCE-hSK (#41814), pCE-hUL (#41855), pCE-mp53DD (#41856); and pCXB-EBNA1 (#41857); all from Addgene) and nucleofection carried out with Amaxa Nucleofector II^TM^. PBMCs were recovered in StemPro34 medium and transferred to 2 wells of a vitronectin-coated 6 well plate containing each 2 ml StemPro34 medium. On the next day (D 1), 1 ml of N2B27 medium supplemented with bFGF (50 ng/ml) and sodium butyrate (NaB, 0.25 mM was added to each well. Until D 7, daily half media change and on D 8 full media change with supplemented N2B27 was performed. On D 9, medium was replaced by Essential 8 Flex and nascent colonies grown for about 3 weeks. Cultures were supplemented with 5 µM ROCK inhibitor before single colonies were picked (n=8-12) under microscopic control using 25 gauge 1 ½ inch needle and 100 µl tips. Remaining colonies were pooled and both singly isolated and pooled colonies were separately expanded on vitronectin-coated plates. Finally, iPSC populations were purified using Tra1-60 microbeads according to manufacturer’s instructions (Miltenyi Biotec^TM^, see also 1.3.2).

### PBMC collection at MPIP

PBMCs were isolated from blood sampling (see above) using SepMate-50 tubes (StemCell Technologies^TM^) according to manufacturer’s instructions. Cell pellets were resuspended in 1-5 ml Erythroid Expansion Medium (EEM)^48^, expanded for reprogramming (see below), or resuspended in EEM containing 10% DMSO (5 x 10^6^ cells/ml) for storage under nitrogen.

### Reprogramming of PBMCs to iPSCs at MPIP

Reprogramming was carried out by modification of a previously published protocol^48^. Briefly, PBMCs were enriched and expanded for erythroid progenitors in EEM across 10-15 days, when either nucleofection was performed or stocks were made (freezing medium 90% FBS/10% DMSO).

Nucleofection was performed with Human CD34+ Nucleofector Kit (Lonza^TM^) and Amaxa Nucleofector II. Erythroid progenitors (2 x 10^6^) were resuspended in 100 µl nucleofection buffer and episomal vectors (4 µg MOS, 4 µg MMK, and 2 µg GBX) **were added**. Following pulse application (program T-016), cells were transferred into a single well of a 12 well plate (D 0) containing 2 ml EEM. At D 2, cells were collected in a 15 ml conical tube and centrifuged at 200 g for 5 minutes (min). Supernatant was carefully descanted, cells resuspended in 1 ml DMEM/F12 supplemented with 10% FBS, and transferred into a single well of a 12 well plate coated with vitronectin (5 µg/ml diluted in DPBS). Plate was sealed with parafilm, centrifuged at 200 g for 30 min at 25 °C, and returned to the incubator. After 24 hr, medium was carefully removed, transferred into a 15 ml conical tube, and centrifuged (300 g for 5 min). Meanwhile, Essential8 (E8) medium (0.5 ml) supplemented with NaB (0.25 mM) was added to the emptied well to prevent drying of already adherent cells. Pelleted cells were resuspended in 0.5 ml supplemented E8 and reseeded in the original well. From D 5 onward, medium was replenished every other day until single cells were grown to sizeable colonies. These were isolated singly or as pool (see above) and expanded onto Matrigel-coated 48 wells with E8 medium supplemented with Revita Cell (E8/RC). Tra-1-60 selection was performed after 3-4 additional passages. Cells were washed once with DPBS, dissociated with accutase for 5 min at 37°C, and centrifuged at 300 g for 5 min. After resuspension in 80 µl E8/RC, 20 µl Tra-1-60-labelled magnetic beads (Miltenyi^TM^) were added and selection of Tra1-60 positive cells conducted according to manufacturer’s instructions. Selected cells were seeded in E8/RC until the next day when medium was replaced by E8 only. Around passage 4-7, iPSCs were adapted to StemMACS iPS Brew XF by culturing with an equal mixture of both media for one passage before replacing by StemMACS iPS Brew XF only. For storage, iPSCs were resuspended in freezing medium (50% FBS, 40% DMEM/F12, and 10% DMSO), transferred in a freezing box to -80°C for 24 hr, and thereafter to liquid nitrogen.

### Reprogramming of fibroblasts to iPSCs

Life cultures of primary human skin fibroblasts from were obtained from Coriell. After overnight recovery, medium was replaced by FBM (DMEM, 10 % FCS, 1x MEM-NEAA) and renewed every 3-4 days. When cells reached 90 % confluency, fibroblasts were washed with DPBS and pretreated at room temperature (RT) with 0.5 mM EDTA (UltraPure™) for 5 min before cells were dissociated with 1x Trypsin/EDTA. Dislocated cells were centrifugated in FBM (300 g for 5 min at RT), reseeded, and expanded for 3-5 passages.

Reprogramming of fibroblasts was conducted by modification of a previously published protocol^49^. Briefly, fibroblasts (2 x 10^6^), freshly plated 24 hr before nucleofection, were resuspended in 100 µl nucleofection buffer (Nucleofector Kit R (VCA-1001, Lonza^TM^). Following addition of 20 µg CoMiP 4in1 plasmid (#63726, Addgene), fibroblasts were nucleofected (Amaxa Nucleofector II, program U-023). Recovered fibroblasts were plated on Matrigel in 2 wells of a 6-well plate (D 0). On the following day, media was replaced by FBM supplemented with 50 µg/ml ascorbic acid and a cocktail of inhibitors (0.3 mM PFT1, 3 mM EPZ 4777, and 0.2 mM NaB). Medium was replenished on D 2 and half-changed to supplemented E7 media on D3. From D 4 onward, supplemented E7 was replaced daily over 5 days. Media was half changed to supplemented E8 media on D 10 and fully replenished with supplemented E8 from D 11 onward until the first iPSCs colonies appeared around D 16 to 25. Only then, supplements were omitted. Colonies were isolated singly or as pool, expanded, and Tra-1-60 selected as described above. Throughout this study, iPSC cell lines derived from pools of 25-45 individual colonies were used.

### Culturing of iPSCs

Frozen iPSCs were thawed in the water bath, collected in DMEM/F12 and centrifuged at 300 g for 5 min. Pellet was resuspended in StemMACS iPS Brew XF supplemented with 100x RevitaCell Supplement and seeded on plates that were coated with Matrigel (diluted 1:100 in DMEM/F12) for at least 30 min. Medium was changed every day by fresh StemMACS iPS Brew XF and cells were passaged twice per week. For passaging, cells were dissociated with gentle cell dissociation reagent for 5 min at 37°C, collected in DMEM/F12, centrifuged at 300 g for 5 min and seeded 1:6 in StemMACS iPS Brew XF on Matrigel-coated plates.

### iPSC characterization: Digital karyotyping

#### DNA extraction and genotyping

In order to characterize the original as well as the generated cell type by digital karyotyping, DNA of PBMCs or iPSCs from each donor was extracted using the QIAamp DNA Blood Mini Kit from Qiagen. To that end, cells were collected in a 15 ml conical tube, pelleted by centrifugation at 300 g for 5 min and placed on ice. Medium was removed, pellet was dissolved in 200 µl DPBS and transferred into a 1.5 ml reaction tube prepared with 20 µl Proteinase K (from the kit). 4 µl RNase (from the kit) was added per tube and suspension was mixed by pipetting up and down. After adding 200 µl AL Buffer and vortexing for 15 seconds (sec), mixture was incubated at 56 °C for 20 min. 200 µl 100% EtOH were added and mixed with the rest by pipetting up and down. Suspension was transferred to a column provided from the kit and vacuum was applied to let the fluid pass through. After washing the membrane with 500 µl AW1 Buffer and another application of vacuum, membrane was washed with 500 µl AW2 Buffer and vacuum was applied another time. Columns were transferred into a collection tube and centrifuged at 13000 g for 1 min. Flow-through was discarded and tubes were centrifuged at 13000 g for 30 sec. After transferring the column into a new 1.5 ml tube, DNA was eluted by adding 80 µl H_2_O and centrifuging at 13000 g for 1 min. Extracted DNA from PBMCs or iPSC cells were hybridized to the Illumina Global Screening Array-24 (GSA).

#### Analysis digital karyotyping

In order to detect de novo copy number abnormalities (CNA) arising in cultured cell lines or general screening for copy number variation status (CNV), we implemented a comprehensive pipeline which includes a developed digital karyotyping method developed^50^ and freely available in BCFtools package^51^ as “cnv” command. In addition, our pipeline (available at https://gitlab.mpcdf.mpg.de/luciat/cnv_detection.git) includes also quality control for genotype and sample calls as well as sample label mismatch detection and correction. The goal is to determine possible differences between cell lines and starting material of derivation in term of copy number variation using genotyping arrays. This genomic screening for chromosomal abnormalities is used as a quality control to establish and maintain stem cell lines. We indicate with line the single probe/sample and with donor the individual from which multiple lines can be derived (e.g. iPSCs and fibroblast).

Following raw data acquisition, Genotyping (GT) module of GenomeStudio software provided by Illumina was used to call genotypes. For each probe, GT module estimates Log R ratio (LRR) and B-allele frequency (BAF) using a clustering module applied to the distribution of signal intensities. Samples quality is evaluated and quality control of genotype calls are curated. In particular, SNPs statistics from GenomeStudio are reviewed and filtered based on the protocols^52^. For autosomal chromosomes and pseudoautosomal regions in chromosomes X and Y, variants with any of the following conditions are excluded: cluster separation ≤ 0.3, AA R Mean ≤ 0.2, AB R Mean ≤ 0.2, BB R Mean ≤ 0.2, AB T mean ≤ 0.1 or > 0.9, Het Excess < -0.9 or > 0.9, MAF > 0 and AB Freq = 0, AA Freq = 1 and AA T Mean > 0.3, AA Freq = 1 and AA T Dev > 0.06, BB Freq = 1 and BB T Mean < 0.7, BB Freq = 1 and BB T Dev > 0.06. Finally, we also excluded variants with call frequency < 0.98 if the total number of probes is higher than 500, otherwise lower or equal than 𝑓(𝑛) = 0.0004 ∗ 𝑛 + 0.7804 with 𝑛 the number of probes in the chip considered, so that a study with a small number of samples is not excessively penalized. For variants in the haploid chromosomes instead, we initially inferred the sex of each sample. If (i) percentage of genotype called as AB in chromosome X < 0.01 and (ii) percentage of not called (NC) genotypes in chromosome Y is < 0.7 then the sample is assigned as “male”. Instead, if both conditions (i) and (ii) are not satisfied the sample is assigned as “female” and if only one of the two conditions is met then the sample is assigned as “undefined”. Afterwards, a variant in chromosome X is excluded if the fraction of AB calls in males with respect to the total number of males is higher than 0.5 and a variant in chromosome Y is excluded if the fraction of assigned calls with R > 0.2 in females with respect to the total number of females is higher than 0.5. In chromosome Y, variants such that the call frequency in males is lower than 0.98 or the function 𝑓 previously defined depending on the number of samples are additionally excluded. The final set of variants is used to compute sample call rates with should be > 0.98 to be considered reliable.

Then, sample mismatches or mislabelling are checked using genotype assignment (AA, AB or BB) and a percentage of matching genotypes for each pair of the available lines is computed as the percentage of variants with the same call. Lines from same donor should show genotype match around 99%. If the match is lower than 97% for the same donor or there is one or more lines that match a certain donor but are not assigned to it, then the matching samples are reassigned with a new name.

Finally, copy number differences between starting material (e.g. PBMC) and derived lines (e.g. iPSC) from the same donor are inferred using a hidden Markov model algorithm implemented in BCFtools (cnv command)^50^. The program is run in the pairwise mode with the default parameters. For the CNV detection of a single line instead, the program is run on single mode based on a single-sample hidden Markov model. CNA calls are then filtered to reduce the number of false calls (see Methods section^53^) and only CNA with the following characteristics are retained: length higher or equal than 0.2 Mb, quality score higher than 2, number of markers and heterozygous markers for copy number 1 and 3 respectively higher than 10.

Genotype imputation, PRS calculation, and CNV analysis

#### Whole genome imputation of human genotyping data

GSA array genotyping data for 184 human samples corresponding to the case/control cohort used in this study was processed by Illumina Genome Studio <> PLINK1.9 and PLINK2 were used for all quality control assessments; first a per SNP and per sample missingness count was performed with parameters set at --maf 0.01 --geno 0.02 --mind 0.02. The dataset was then filtered for sample repeats, identity by descent was used to identify close relatives which were temporarily removed (PI HAT threshold = 0.0625) to assess population stratification and identify population outliers. The latter was done by first pruning the SNPs by --geno 0.02 --hwe 1e-3 --indep-pairwise 200 100 0.2 --maf 0.05 —set-hh-missing and excluding the major histocompatibility complex (MHC) as well as the interstitial inverted duplication INV8, this was followed by creating an multidimensional scaling (MDS) clustering using the following parameters: —cluster --mds-plot 10 eigendecomp. Outliers were discarded using a threshold of 4 standard deviations and relatives were added back to the dataset. SNPs were evaluated and filtered based on Hardy-Weinberg equilibrium with a threshold of 1e-6. SNP names were updated to match those of the 1000 Genomes Phase 3 which was the reference panel used for imputing the samples. Duplicated SNPs were removed and remaining SNPs were checked and strand orientation corrected if needed. The dataset was phased using shapeit v2.r837 and windows of 5Mb were imputed using IMPUTE version 2.3.2 using the parameters -pgs_miss -filt_rules_l -buffer 500. Post-imputation quality control was applied by identifying empty or low SNP count blocks which were either discarded or reimputed by moving the window location and/or size. In order to filter high quality imputed SNPs, an info score great than 0.8 and minor allele frequency threshold of 0.01 were used to select the final SNPs.

#### PRS calculation

We calculated PRS for schizophrenia based on prior GWAS of schizophrenia using the PGC3 release^3^, Mullins et al. for BD^6^ and Wray et al. for MDD^54^. PRS-CS^55^ was used to infer posterior SNP effect sizes under continuous shrinkage priors and estimate the global shrinkage parameter (φ) using a fully Bayesian approach with genotype dosage data.

#### CNV analysis

Bipolar disorder, Major Depressive disorder and Autism associated Copy Number Variations (CNV) were obtained from Zarrei et. al.^56^, only CNVs labeled as high confidence were used. CNVs were called in all our samples and later compared to the known disease associated CNVs mentioned above by overlapping their genomic coordinates, this was done using bedtools intersect v2.26.0, a match was only called when a minimum overlap of 90% of both CNVs being compared was covered. The same overlapping procedure was applied to SCZ CNVs obtained from Marshall et al.^57^, which were first lifted over from hg18 to hg19 genome assembly using UCSC^58^ liftover tool (http://genome.ucsc.edu/cgi-bin/hgLiftOver).

#### iPSC characterization: Immunocytochemistry (ICC)

Initially, attached cells were washed 1x with DPBS. DPBS was removed and fixation was performed with 4% PFA in DPBS for 10 min at RT. After removal of fixation solution, 3x washing steps with DPBS were performed. Samples were stored up to 2 weeks at 4°C for subsequent steps. After permeabilization with 0.1% Trition-X-100 in PBS for 10 min at RT blocking was performed with blocking solution of 0.1% Trition-X-100 and 1% BSA in PBS for 60 min at RT. Primary antibodies were added in blocking solution and incubated overnight at 4°C. At the next day, primary antibody solution was removed with 3x washing with PBS. Respective secondary antibodies in PBS were added and incubated for 60 min at RT. Secondary antibody solution was removed with 3x washing with PBS, Finally, samples were washed 1x with water and mounted onto microscope slides. If mounting medium did not contain DAPI, nuclei counterstaining was performed with DAPI, supplemented during the 2nd washing step. At the LMU Munich, imaging was performed on an Axio Observer.Z1 (Zeiss) inverted microscope and image analysis was performed with ZEN (Version 3.3, Zeiss) and Fiji^59^.

### iPSC characterization: Embryoid body (three germ layer) differentiation

#### Embryoid body production

In order to confirm the differentiation potential into all three germ layers, embryoid body (EB) formation was performed for each reprogrammed cell line^60^. To that end, iPSCs were seeded in iPS Brew medium in one well of a 6well plate coated with Matrigel and grown until 90 % confluency. Medium was aspirated and cells were washed once with 1 ml DPBS before being dissociated by 1 ml Gentle Cell Dissociation Reagent for 5 min at RT. 1 ml iPS Brew was added and cell suspension was transferred into a 15 ml conical tube without disrupting the dissociated colonies. After centrifugation at 300 g for 5 min, pellet was solved in 5 ml EB medium consisting of Knockout-DMEM (88%), Knockout Serum Replacement (10%), Non-essential amino acids (1%) and L-Glutamine (1%) by pipetting up and down maximally 3 times, transferred into a 60 mm dish and incubated at 37°C. Medium was changed on Day 2 and Day 4 by transferring the EB suspension gently into a 15 ml conical tube and adding 2 ml EB medium into the dish in order to avoid drying of remaining EBs. After letting EBs sediment in the conical tubes for 10-15 min, supernatant was aspirated, EBS resuspended with 3 ml fresh EB medium and transferred back into the 60 mm dish. While changing the medium on Day 5 in the same way as before for 4 ml of EB suspension, remaining 1 ml suspension was transferred into a separate 15 ml conical tube, cells were sedimented separately and resuspended in 2 ml EB medium for being plated into two wells of a 24well plate coated with Matrigel (1ml/well). Medium was changed every other day until Day 12-15. For harvesting the cells in the dish, EBs were transferred gently into 15 ml conical tube and centrifuged at 300 g for 5 min. After removing supernatant, pellet was frozen at -80°C. Attached cells in the 24well plate were fixed in 16% Paraformaldehyde and were stored at 4°C until being stained.

#### RNA-Seq library production (from EBs)

QuantSeq 3’ mRNA-Seq Library Prep Kits (Lexogen) was used to prepare Illumina sequencing libraries. Based on manufacturer’s protocol 500 ng of RNA from each sample was used to prepare the RNA-Seq libraries. Library’s size distribution and quality were quantified by high-sensitivity DNA chip (Agilent Technologies).

#### Characterization of germ layer differentiation potential

In order to assess iPSC differentiation potential into cells from the three embryonic germ layers ectoderm, mesoderm and endoderm, we followed a similar strategy previously implemented in the iPSC Scorecard concept and its derivatives^60^, relying on 15 days long random differentiation of iPSCs into embryoid bodies, followed by bulk RNA-Seq of the respective EBs. In addition, we generated representative RNA-Seq profiles from a subset of the iPSC lines. Subsequently, we used established markers for pluripotent, ectodermal, mesodermal and endodermal cells and computed the “differentiation potential” of each iPSC line into these germ layers based on the RNA-Seq profiles of the EBs, following the same approach as implemented by Tsankov et al.^61^. Briefly, to analyze the germ layer marker gene expression in each EB sample, we computed the one-sided t-statistic comparing the gene’s expression against its distribution across the iPSC lines. As negative control, we also computed the one-sided t-statistic for each marker in each iPSC line compared to its expression in all other iPSC lines. In addition, we computed the log_2_ fold change of each marker gene between the average EB and average iPSC expression level as weight for each marker gene and calculated the germ layer differentiation potential based on the marker gene sets for each germ layer, analogous to the approach employed by Tsankov et al.^61^ and display the results in Extended Data Fig. 1k,l.

### Differentiation of excitatory cortical neurons from iPSCs

In order to investigate potential differences in neurons derived from patients with mental illness and healthy controls, iPSC from 106 different donors were differentiated into induced cortical glutamatergic neurons (iNs) according to previous protocols^26,62^ by using doxycycline-inducible overexpression of the transcription factor NGN2 and cultured those neurons, together with murine astrocytes after seven days, for a period of 49 days.

#### Lentivirus production

Human embryonic kidney cell line HEK293 was cultured in DMEM (high glucose, pyruvate) supplemented with 10% FBS. Around 80 % confluency, cells were dissociated with 2x Trypsin and collected in DMEM/10% FBS. After centrifugation at 300 g for 5 min and counting with Neubauer Chamber, 5 x 10^6^ cells were seeded in 8 ml DMEM/10% FBS into a cell culture dish (8.7 cm^2^) and incubated for 20-24 hours at 37°C. Cells were washed once with 5 ml DMEM/F12 and afterwards cultured in 8 ml DMEM/F12. Two mixes forming the plasmid mix and the transfection reagent mix were prepared. For each cell culture dish 6 µg psPAX2, 3 µg pMD2.G (VSVG) and 9 µg pTet-O_Ngn2-puro or FUW-M2rtTA were prepared in 0.5 ml DMEM/F12 forming the plasmid mix. In a second mix 54 µl transfection reagent LipoD293 DNA was added in 0.5 ml DMEM/F12. After vortexing both mixes briefly, LipoD293 DNA mix was added to the plasmid mix, vortexed briefly again and incubated for ten min at RT. After adding 1 ml transfection mix dropwise to the cells and incubating for 18 hours at 37 °C, medium was replaced by 20 ml DMEM/10% FBS and incubated for further 48 hours at 37 °C. Medium was collected, filtered through 0.45 µm sterile Acrodisc syringe filter in a 50 ml conical tube and supplemented with 5 ml Peg-it Virus Precipitation Solution. After inverting the tubes five to ten times, mixture of viral particles and Peg-it Virus Precipitation Solution was incubated for 72 hours at 4 °C. After incubation time, mixture was centrifuged at 1500 g for 30 min at 4°C. The supernatant was discarded, pellet with precipitated viral particles resuspended in 250 µl cold DMEM/F12 (1:100), aliquoted on ice and stored at -80 °C.

#### Mouse astrocyte preparation

25-35 P1 mice were sacrificed in each batch and cerebral cortices were isolated in PBS supplemented with 2.5% FBS. Dissected cortices were transferred into a 15 ml conical tube and centrifuged briefly to remove the remaining PBS/10% FCS. After washing the pellet twice with 5 ml PBS, 2 ml 1% DNAse in PBS was added and incubated for 5 min at 37°C in the water bath followed by adding 2 ml 2.5x trypsin and further incubation for 5 min at 37°C. Enhancing enzymatic dissociation, cell suspension was pipetted up and down 15 times starting with a 10 ml serological pipette, incubating for 5 min at 37 °C, pipetting up and down 15 times with a 5 ml serological pipette, incubating for 7 min at 37 °C, ending in pipetting up and down 10 times with the 1 ml filter tip. After dissociation single cells were filtered through a 100 µm filter mesh and collected in a 50 ml conical tube containing DMEM (high glucose, pyruvate) supplemented with 10% horse serum (HS). Filter mesh was washed twice with DMEM/10% HS and cell suspension was centrifuged for 10 min at 300g. Supernatant was discarded carefully, cell pellet was resuspended in 5 ml DMEM/10% HS and filtered through a 40 µm filter mesh into a new 50 ml conical tube containing DMEM/10% HS. Filter mesh was washed twice with DMEM/10% HS and cells were seeded into four 75 cm^2^ flasks that were coated with Attachment Factor Protein (1x) for at least 30 min at 37°C. Medium was changed to fresh DMEM/10% HS after two days and subsequently twice per week. Astrocytes were passaged after seven to nine days by incubating the cells with 2x Trypsin for 10 min at 37°C, collecting in DMEM/10% HS, centrifuging at 300 g for 5 min and seeding the cells 1:2 to 1:2.5 on Attachment Factor-coated flasks. To end up always in the same passage number (P.3), astrocytes were either passaged another time after seven more days or were frozen transiently. To that end, cells were dissociated with 2x Trypsin, collected in DMEM/10% HS, centrifuged at 300 g for 5 min and either frozen in 90% DMEM/10% HS combined with 10% DMSO or seeded again 1:2 to 1:2.5 in DMEM/10% HS on Attachment Factor Protein-coated flasks.

#### Neuronal differentiation

At Day 0, 10cm dishes were coated with Matrigel (Matrigel in DMEM/F12; 5 ml/ 10cm dish for at least 1 hour at 37°C. iPSCs were separated with Accutase for 5 min at 37°C, diluted 1:1 in DMEM/F12 supplemented with 1% BSA, centrifuged at 300g for 5 min and resuspended in StemMACS iPS Brew XF supplemented with Polybrene (6µg/ml) and 1x RevitaCell. Cells were counted, cell suspension was adjusted to a concentration of 500,000 cells/ml) and lentiviral infection with pTet-O_Ngn2-puro and FUW-M2rtTA was performed. After incubation for 10 min at RT, subsequent seeding (25,000 cells/cm²) on Matrigel-coated plates was performed.

At the next day (Day 1), media was replaced by KSR media, which consists of KO DMEM with 15% Knockout Serum Replacement, 1x NEAA, 1x Glutamax and 50 µM beta-mercaptoethanol, supplemented with 10 µM SB431542, 2 µM XAV939, 0.1 µM LDN-193189 and 2 µg/ml doxycycline to start neuronal patterning and directed differentiation. At Day 2, KSR media replaced by a 1:1 mixture of KSR media and N2 media, which consists of DMEM/F-12, 1x Glutamax, 3mg/ml glucose and 1x N-2, supplemented with 5 µM SB431542, 1 µM XAV939, 0.05 µM LDN-193189, doxycycline and 10 µg/ml puromycin to select for successfully co-infected cells. At Day 3, media was replaced by N2 media supplemented with doxycycline and a final day with puromycin.

At the following day (Day 4), young iNs were dissociated by treatment with Accutase for 5 min at 37°C. After diluting cell suspension in Accutase in doubled volume of 1%BSA in DMEM/F12, cells were triturated by pipetting up and down 3x with an 1ml tip. After centrifugation, cells were seeded with 60.000 cells/cm² (200,000 cells/glass coverslip in a 24well for Patch Clamp Recording, 150.000 cells/well for MEA recordings) in NBM media, which consists of Neurobasal medium, 1x Glutamax, 1x NEAA, 1xB27 without vitamin A, 3 mg/ml glucose, 2% fetal bovine serum (FBS), freshly supplemented with 10 ng/ml BDNF, 10 ng/ml CNTF, 10 ng/ml GDNF and 2 µg/ml doxycycline, on dishes that were coated with Poly-L-Ornithine (15 µg/ml) overnight at 37°C and after 3x washing with DPBS at least 6 hours at 37°C with Laminin (1µg/ml) and Fibronectin (2 µg/ml). Regarding functional assays as patch clamp and MEA recordings, glass coverslips or glass plates were coated with Poly-L-Ornithine (20 µg/ml) in borate buffer (100 mM boric acid, 25mM Sodium Tetraborate in deionized H_2_O) for 6 hours at 37°C, Wells were washed 3x with H_2_O (WFI), dried overnight at RT and coated the next day with Laminin (10 µg/ ml) and Fibronectin (20 µg/ ml) for at least 6 hours at 37°C.

At Day 7, murine astrocytes were added by full media change with a density of 5000 mouse astrocytes/cm² on the iNs. After three days, last full media change was performed and 4 µM AraC was added to NBM media to stop glial cell division. Subsequently, half media changes were performed twice a week. Doxycycline was removed from NBM media after 21 days and replaced by 1µg/ml laminin. At Day 24, NBM media was replaced by BrainPhys media, which consists of BrainPhys Neuronal Medium, 1x Glutamax, 1x NEAA, 1xB27 without vitamin A, 3 mg/ml glucose, 3% fetal calf serum (FCS), freshly supplemented with 10 ng/ml BDNF, 10 ng/ml CNTF, 10 ng/ml GDNF and 1µg/ml laminin to foster neuronal maturation. At Day 49 experiments for endophenotyping were performed.

### Endophenotyping of iPSC-derived neurons

Investigating differential expression of genes between iNs derived from individuals with schizophrenia (ISCZ) or bipolar disorder (BD) compared to iNs derived from Ctrls, various experimental approaches were performed.

### BulkRNAseq – preparation

#### RNA sample collection

iNs were collected after 49 days of differentiation and RNA was isolated by using the RNA Clean and Concentrator-5 Kit (Zymo Research). Whole network was detached and collected in 1.4 ml cold DPBS in a 1.5 ml reaction tube. Cells were centrifuged at 300 g for 10 min at 4°C. PBS was removed and 500 µl QIAzol Lysis Reagent (Qiagen) was added per tube. Sample was incubated for 10 min at RT and subsequently either stored at -80 °C or further processed directly. After adding 100 µl chloroform, sample was vortexed, incubated for 2 min at RT and afterwards centrifuged at 10000 g for 15 min at 4 °C. Upper phase was collected in a new 1.5 ml reaction tube and same volume 100% ethanol was added and mixed by pipetting up and down. Sample was loaded on the Zymo-column provided by the kit and centrifuged for 30 sec at 10000 g. After adding 400 µl RNA Prep Buffer and further centrifugation for 30 min at 10000 g, sample was washed with 700 µl RNA Wash Buffer and centrifuged for 1 min. Flow through was discarded and sample was centrifuged again for 30 min at 10000 g to remove residual Wash Buffer. After transferring column into new 1.5 ml collection tube, RNA was eluted by adding 15 µl DNA/RNAse free water and centrifuging for 1 min at 13000 g. In order to increase the outcome, the last step was repeated by adding eluted 15 µl again on column and centrifuging again for 1 min at 13000 g.

RNA was treated with DNAse using DNA-*free^TM^* DNA Removal Kit (Invitrogen^®^). Therefore 8 µl RNA was mixed with 1 µl 10x DNase I Buffer and 1 µl recombinant DNase I (rDNase I), and incubated for 30 min at 37 °C. 2 µl DNAse Removal Buffer was added and mixed with sample, and further incubated for 2 min at RT. The inactivation beads were spined down by centrifugation for 30”, supernatant was transferred into a new tube and RNA was quantified by using RNA ScreenTape on 2200 TapeStation from Agilent Technologies and assessed with 2200 TapeStation Controller Software. For that purpose, 5 µl RNA Buffer were mixed with 1 µl RNA sample by vortexing for 1 min. Sample was centrifuged briefly, incubated for 3 min at 72 °C and subsequently incubated for 2 min on ice before being loaded on the TapeStation. Samples with a RIN above 8 were further processed.

#### RNA-Seq library preparation

Full length: RNA was retrotranscribe and amplified using the SMART-Seq v4 Ultra Low Input RNA Kit for Sequencing (Clonotech). In particular, 2 ng of high quality RNA was mix with 0.3 µl 10x Reaction buffer (prepared following the manufacturer’s instructions) and 0.5 µl SMART-Seq CDS Primer II A. RNA secondary structures were destabilized at 72°C for 3 min. Afterwards, a mix containing 1 µl 5x First-Strand Buffer, 0.3 µl SMART-seq v4 Oligo, 0.2 µl RNase Inhibitors and 0.5 µl SMARTScribe enzyme was added to each reaction. Reverse transcription was was performed at 42°C for 90 min and the reverse transcriptase was inactivated by 10 min at 70°C. For the cDNA amplification step, to the last reaction a mix of 7.7 µl containing 6.3 µl 2x SeqAmp Buffer, 0.3 µl PCR Primer IIA and 0.3 µl SeqAmp Polymerase was added.

The amplified double-strand cDNA was purified using 1.0X AMPure XP beads (Beckman Coulter), washed twice with 80% EtOH and eluate in 13 µl Elution Buffer (from Clonotech kit) and quantified on BioAnalyzer, (Agilent) using the High Sensitivity DNA chip.

Forty picograms of the purified double-strand cDNA were fragmented using the Nextera XT DNA Library Preparation Kit (Illumina). The reaction was carried out in 5.1 µl adding to the appropriate volume of sample 2.5 µl Tagment DNA Buffer and 1.3 µl Amplicon Tagment Mix. The reaction was incubated at 55°C for 5 min and rapidly cooled down at 10°C. The tagmentation was inactivated incubating the samples at room temperature for 5 min with 1.3 µl Neutralize Tagment buffer.

A different combination of index oligo from the Nextera XT Index Kit v2 Set A (Illumina) was used for each sample and amplified in a 12.8 µl reaction using 3.8 µl Nextera PCR Master Mix. Indexed libraries were purified using 1.8X AMPure XP beads (Beckman Coulter), washed twice with 80% EtOH and eluate in 23 µl.

The final libraries were quantified on the BioAnalizer using the High Sensitivity DNA chip, multiplexed in equimolar pools and sequenced on HiSeq4000 (Illumina).

### BulkRNAseq – analysis

#### RNA-Seq data processing and analysis

Paired-End raw RNA-Seq read sequences were aligned against the GRCh38 genome assembly with the GENCODE version 27 transcriptome definition using the STAR aligner^63^ version 2.7.3a with the following parameters: --outFilterType BySJout --outFilterMultimapNmax 20 -- outFilterMismatchNmax 999 --outFilterMismatchNoverReadLmax 0.04 --alignIntronMin 20 -- alignIntronMax 1000000 --alignMatesGapMax 1000000 --alignSJoverhangMin 8 -- alignSJDBoverhangMin 1 --sjdbScore 1 --readFilesCommand zcat --outSAMtype BAM Unsorted SortedByCoordinate --quantMode GeneCounts --outSAMstrandField intronMotif. Subsequently, duplicates were marked and library quality metric were computed using picard tools (http://broadinstitute.github.io/picard/). In addition, reads were also aligned to GRCh37 and GENCODE version 27 for detailed comparison analysis with human post mortem data and eQTL calling, which was performed on GRCh37. Following quality control, gene level read counts were obtained using the Rsubread package, collapsing exon level reads onto gene level features based on GENCODE version 27 with the following parameters: countMultiMappingReads=FALSE,ignoreDup=FALSE,sGTFAnnotationFile=TRUE, isPairedEnd=T. Quantitative expression analysis was performed with DESeq2 version 1.22.2. Following size factor estimation, we excluded genes with total read counts across all samples below the 20^th^ and above the 99^th^ percentile as well as mitochondrial genes. Finally, we performed standard differential expression analysis testing for effect of diagnosis using differentiation site and sex as covariate including only one RNA-Seq sample per donor. Genes showing an adjusted p-value :<0.01 and a minimal absolute log_2_ fold-change ;: 0.5 between Ctrl and SCZ or Ctrl and BD (adjusted p-value :<0.05) were defined as differential and used in subsequent analysis. To account for potential technical confounders originating from the library preparation etc., we performed PCA on the following technical parameters determined for each library using picard tools: PF_BASES, PERCENT_DUPLICATION, PCT_CODING_BASES, PCT_UTR_BASES, PCT_INTRONIC_BASES, PCT_INTERGENIC_BASES, MEDIAN_5PRIME_BIAS, MEDIAN_3PRIME_BIAS. PF_BASES was log2 transformed prior to PCA. We then included the first principal component (TechnicalPC1) from this analysis as covariate in all downstream analyses.

#### Identification of novel polyadenylation sites in the 3’UTR in iNs

To identify alternative polyadenylation sites and quantify differential polyadenylation site usage, we followed the recommendations from a recent benchmarking study comparing different experimental and computational approaches^40^. To that end, we first detected all polyadenylation sites (PAS) in use in iNs using 3’RNA-Seq in iNs at day 49 from 6 distinct donors^64^ by mapping their 3’UTRs. We then detected all polyadenylated transcribed regions within the union of these RNA-Seq libraries using the R package transcriptR^65^ in the following manner: We built a reference collection of candidate 3’transcript fragments using the constructTDS function with fragment size set to 250 and the swap.strand option set. Next, we overlapped the transcript fragment collection with all genes from Gencode version 27 and estimated the background distribution of detected fragments across the genome using the estimateBackground function with an FDR cutoff of 0.01. Subsequently, we identify polyadenylated transcripts using the detectTranscript function with a gap distance of 75, followed by merging neighboring candidate transcripts fragments within 100bp of each other. Finally, we only retain transcripts with a minimum size of 50bp, a maximum size of 2kb, supported by at least 10 independent sequenced fragments, an FPKM of greater than 3 and overlapping with a known Gencode gene. The resulting transcript collection represents transcript fragments detected based on their polyadenylated 3’-end region. Finally, we use the APA analysis software QAPA to identify those transcripts with end-coordinates located within or beyond known 3’UTRs of Gencode genes using the *qapa build* command. The 3’UTR end-coordinates correspond to novel and known PAS sites in iNs. All internal PAS are not considered further. We next defined coordinates of all individual 3’UTRs based on the coding sequence end-coordinate of each transcript and the PAS end-coordinate for each detected individual transcript fragment.

Given that many Gencode transcripts harbored more than two PAS within a single known 3’UTR, we next reduced this complexity to two PAS sites/UTR variants for each transcript, selecting the most distal PAS and the dominant proximal PAS. To that end, we segmented the entire 3’UTR region of each gene, extending from the CDS end-coordinate to the most distal PAS, into non-overlapping 3’UTR fragments based on all PAS located within the UTR. Subsequently, we quantified the number of overlapping read fragments from our full-length RNA-Seq data in iNs across 100 RNA-Seq samples and defined the UTR fragment with the highest mean normalized read count as the dominant proximal UTR/PAS site. These analyses gave rise to a maximum of two 3’UTR regions for each gene that were used for all subsequent analyses (**Supplementary Table 3**, APA sites). For differential APA analysis, we next quantified the normalized read count in all full-length RNA-Seq samples for each of the non-overlapping 3’UTR regions for each gene, counting all reads overlapping with the proximal or distal UTR region respectively. From these values, r_prox_ and r_dist_, we computed the percentage of distal 3’UTR used for each gene *j* and each iN donor *I*, analogous to previous studies^66,67^ as R_ij_= r_distij_/(r_proxij_+r_distij_). To quantify differential PAS site usage between HC and SCZ in iNs, we employed a beta regression approach as reported previously using a logit link function for the mean model, the identity function for the precision model as well as a maximum likelihood estimator^66^, correcting for gender, batch and TechnicalPC1 (see the RNA-Seq analysis), testing for diagnosis status. Similarly, we also corrected the R_ij_ values for gender, batch and TechnicalPC1 using beta regression and used the residuals to compute the average difference in R values between the groups (ΔAPA, **Supplementary Table 3**). P-value were adjusted using the q-value method^68^ and filtered based a q-value :<0.05.

#### *3’APA* analysis of iNs RNA-Seq and post mortem RNA-Seq data

Analysis of differential APA in human postmortem samples was performed in a similar manner to dAPA analysis, quantifying normalized read counts across all CommondMind RNA-Seq libraires for samples with HC or SCZ status using the same 3’UTR library that was employed in iNs. dAPA analysis was conducted as before using beta regression adjusting for the following covariates: institution, gender, age of death, post mortem interval, RIN, RIN2, clustLIB, the first 5 genotype PCs capturing ancestry as well as the first 3 TechnicalPCs derived from the PCA of all RNA-Seq library quality parameter of the corresonding libraries. All dAPA events with a q-value :<0.1 were considered significant.

#### Association analysis APA-PRS

To enable gene expression or APA association analysis with PRS, we first computed a cumulative measure of gene expression or APA across all differentially expressed or polyadenylated genes between SCZ and HC or MDD/BD and HC by averaging over their z-score normalized gene expression or R values for each sample separately. Following outlier filtering using Tukey’s criterion for the averaged values, we performed association testing using a linear model to estimate the association between cumulated differential gene expression or cumulated dAPA with PRS for SCZ, BD and MDD separately.

#### LD-score regression and partition heritability analysis of transcripts subject to dAPA

To estimate whether or not genomic loci with transcripts that are subject to APA are enriched for psychiatric disease associated heritability, we performed LDscore regression and partition heritability analysis as previously described^43^. To that end, UTR regions extended by xx subject to APA in iNs or human postmortem PFC were intersected with the LDSC annotation files using bedtools^69^ Version: v2.27.1-1-gb87c465. and formatted as input to the S-LDSC software^70^. LD scores were then calculated separately for each category (using LD Score Regression (LDSC) Version 1.0.1). Partition heritability analysis was performed according to current recommendation using the baseline model version 1.2^43^. For all annotation categories, a cell-type-specific analysis was performed using the *--h2-cts* flag to estimate disease association for the following traits: Major Depressive Disorder, Antidepressant treatment resistant, Any psychotic experience, Schizophrenia, Ever Smoked, Autism Spectrum Disorder, Bipolar Disorder, Major Depression, Psychotic Experience, Coronary Artery, Disease, Height, Type 1 Diabetes, Type 2 Diabetes, Neuroticism, LDL, Celiac, Years of Education. This was carried out using each disease’s corresponding publicly available GWAS summary statistics obtained from https://alkesgroup.broadinstitute.org/LDSCORE/all_sumstats/.

Finally, we report the cell and condition specific enrichment of all by traits P-value of tau and z-scores based on the regression coefficient and coefficient standard error.

#### Transcript Pathway analysis

In order to annotate transcripts subject to APA in iNs from ISCZ, we performed pathway and ontology enrichment analysis of this gene set using the enrichR meta-database^71^ and SYNGO^46^.

#### Power analysis

For detailed power analysis, we utilized the R-framework RnaSeqSampleSize^72^, relying on a negative binomial model to simulate read count distributions from individual sample groups. We followed the standard protocol for power estimation based on RNA-Seq samples from the Ctrl and SCZ sample group, including only one sample per donor and using the same samples employed for the differential expression analysis in the main manuscript. Subsequently, genes were filtered based on raw read counts including only genes with at least 11 reads (20^th^ percentile) and less than 478780.4 (99^th^ percentile) total reads per gene across the entire dataset. Power estimation analysis was conducted based the remaining gene set. Subsequently, we used the “est_count_dispersion” function of the RnaSeqSampleSize package with a 2 group design to estimate the dispersion, resulting in disp = 0.28525. We rounded this to the nearest digit and subsequently worked with a dispersion estimate of 0.3 We then used this estimate and a maximal sample size of 60 samples per group to compute power curves for different fold-change and FDR combinations using the “est_power_curve” function with lambda0 = 200, phi0 = 0.3, n = 60, f = 0.1 or 0.05 or 0.01 and rho = 1.5 or rho=2. The results are shown in **Extended Data Fig. 3d**.

### Single-cell RNA-Seq from iNs

#### Sample preparation and sequencing

Multiplexed scRNA-Seq profling involved iNs from 4 distinct donors per library. Neuronal networks were dissociated 49 days after differentiation with papain (10 units/ml) and DNAse (100 units/ml) diluted in DMEM/F12 for 10 min at 37 °C to facilitate generation of a single cell suspension. Afterwards, the network was mechanically sheared with a P1000 pipette tip. DMEM/F12 supplemented with 20% FCS was added for centrifugation at 400 g for 5 min at RT. Cells were resuspended in DMEM/F12 supplemented with 1% BSA and 1x RC and pelleted for a final resuspension in 1 % BSA in DPBS supplemented with RNAse Inhibitor, 1x RC, 3 mM Mg(Ac)_2_, and 5 mM CaCl_2_. After removal of network debris with filters, cells were counted. 12,000 cells (3,000 per donor) were loaded on the 10X Chromium machine (10X Genomics). scRNASeq libraries were prepared following manufacturer’s instructions of the Chromium™ Single Cell 3’ Library & Gel Bead Kit v3 (10X Genomics). The scRNA-Seq library was subsequently sequenced/processsed on a NovaSeq 6000 S1 flowcell.

#### Data processing and analysis

For scRNA-Seq read alignment, pre-build GRCh37 genome reference files for 10X cellranger were downloaded from https://www.10xgenomics.com/. Subsequently, cellranger count (version 3.1.0) was used for raw data processing giving rise to feature count matrices. In order to deconvolute the distinct cells from individual donors, we utilized demuxlet supplemented with the relevant donor genotypes to assign the most likely genotype of origin to each cell, following the standard protocol for this approach^73^. More specifically, we created a vcf containing relevant SNPs from the 3’UTR of hg19 transcripts using cftools (version 0.1.17) and the gencode 3’UTR hg19 BED file (downloaded 09.02.2021 from http://genome.ucsc.edu/cgi-bin/hgTables). The resulting vcf encompassed ∼ 98,000 sites. The genotype deconvolution was conducted in demuxlet setting the parameters to demuxlet --alpha 0.0 --alpha 0.0 --vcf vcf_ips_christine_hg19.filtered.recode.sorted.vcf --field GT.

Subsequently, scRNA-Seq data analysis was continued in R (R version 3.6.1)^74^ and Seurat package version 3.2.2 ^75^. A Seurat object was created from the aggregated filtered count matrix, keeping cells with at least 800 and less than 10000 detected features as well as less than 15% mitochondrial RNA reads.

Subsequently, count correction, regression of mitochondrial content, log normalization and pearson residuals were computed according to Seurats SCTransform. PCA on 3,000 most variable features and elbow plot analysis indicated the first 20 principal components to be relevant. These components were then selected for UMAP (min_dist=0.2), nearest neighbour graph construction and clustering analysis (resolution = 0.3, nn.eps = 0.5) using Seurat standard workflow, during which 7 clusters were identified. Subsequently, marker genes were plotted to identify cellular identities (**Extended Data Fig. 2d-f**).

#### Deconvolution analysis of RNA-Seq profiles of iNs

Marker gene sets for deconvolution analysis were defined with Seurat (version 3.2.2)^75^ using the iN scRNA-Seq dataset including all 7 clusters and all donors. The RNA assay of the scRNA-Seq count object was log normalized, scaled and the percentage of mitochondrial gene expression was regressed out. FindAllMarkers was run in order to identify representative genes for each of the 7 clusters setting the following parameters: assay = “RNA”, slot = “data”, test.use = “MAST”, only.pos = TRUE, return.thresh = 0.01. 2,143 hits were significant at adjusted p value ≤0.01, which included 1,875 genes in total. The cluster specificity tau was calculated for each gene^76^ and the resulting sets of markers were filtered to keep all hits with average logFC > 0.35 and tau > 0.6. Consequently, 214 hits covering 210 different genes were used for the deconvolution analysis of 115 RNA-Seq samples of iPSC derived neurons. The deconvolution was implemented using CIBERSORT based R package bseqsc (version 1.0). The bulk counts were CPM normalized and the base matrix of reference gene expression profiles was built on the identified marker genes and the iN scRNA-Seq expression counts. The cluster proportions for the bulk samples were estimated running bseqsc_proportions on the CPM normalized counts and the base matrix. The distribution of resulting cluster proportions for all samples in each diagnoses group was then plotted in **Extended Data Fig. 2g**.

#### Comparison of iN scRNA-Seq profiles to human and murine primary cortical snRNA-Seq

In order to determine the cortical neuronal subtypes and developmental stage that were most closely related to iN cellular identity, we compared previously published^29^ scRNA-Seq data from the developing mouse cortex at developmental stage P1, P7 and P21. For similarity analysis, we first determined all genes expressed in 5% or more of all cells in iNs and mouse samples.

Subsequently, we mapped all murine genes to their human orthologs using the ensemble database. We then aggregated all raw counts per cell identity/cluster and performed VST normalization on the pseudo-bulk samples followed by pearson correlation analysis and multidimensional scaling. Since the first two dimensions primarily captured technical variables (e.g. species and lab of origin), we retained dimension 3 and 4 that separated samples by time points and cellular identity.

### miRNAseq

In order to study miRNA-dependent regulation of genes, miRNAseq was performed.

#### Sample collection

Samples were collected and prepared following the same procedure as for the RNAseq. Isolated and DNAse treated RNA was further processed at Telethon Institute of Genetics and Medicine.

#### Library preparation

Total RNA was quantified using the Qubit 2.0 fluorimetric Assay (Thermo Fisher Scientific) and sample integrity, based on the RIN (RNA integrity number), was assessed using an RNA ScreenTape assay on TapeStation 4200 (Agilent Technologies).

Libraries were prepared from 250 ng of total RNA using the smallRNAseq sequencing service (Next Generation Diagnostics srl) which included, library preparation, quality assessment and sequencing on a NovaSeq 6000 sequencing system using a single-end, 50 cycle strategy (Illumina Inc.).

#### Data processing

The raw data were analyzed by Next Generation Diagnostics srl proprietary SmallRNA-seq pipeline (v 1.0)^77^. Illumina NovaSeq 6000 base call (BCL) files were converted in fastq file through bcl2fastq. Then, trimming and cleaning with Trimgalore (https://github.com/FelixKrueger/TrimGalore) was followed by alignment on the hg19 reference genome using Bowtie^78^. Detection counts were determined with SAMtools^79^. The miRNA genome was build using miRBase v22. Finally, exported tab-delimited text files included the raw counts for subsequent analysis.

#### Analysis

microRNA-Seq data was analyzed using with DESeq2 version 1.22.2. Following size factor estimation, we excluded microRNAs with total read counts less than 100 across all samples. Subsequently, we performed standard differential expression analysis testing for effect of diagnosis using differentiation site and sex as covariate including only one microRNA sample per donor. microRNAs below an adjusted p-value ≤0.05 between Ctrl and SCZ were defined as differential and used in subsequent analysis.

### ATAC-Seq

Chromatin structure was assessed by the assay for transposase-accessible chromatin (ATAC).

#### Sample collection and library preparation

iNs were dissociated at D 49 with 135 U Papain and 1,180 U Deoxyribonuclease I (Worthington) dissolved in 7.5 ml DMEM/ F12 using 1 ml per 6well for 10 min at 37°C. After enhancing enzymatic dissociation by pipetting up and down, cell suspension was collected in DMEM/F12 supplemented with 10% FBS and RevitaCell Supplement (1:100). After centrifugation at 300 g for 5 min, each pellet was resuspended in 1 ml DMEM/F12/10% FBS/RC and counted with Neubauer Chamber. 7 x 10^4^ cells were centrifuged at 500 g for 5 min at 4 °C and supernatant was discarded carefully. After resuspending pellet in cold ATAC-Resuspension Buffer I (RSB I) consisting of 1M Tris-HCl (pH 7.4), 5M NaCl and 1M MgCl_2_ in sterile H_2_O containing 0.1% NP40, 0.1% Tween-20 and 0.01% Digitonin by pipetting up and down 3 times, suspension was incubated for 3 min on ice. 1 ml cold ATAC-RSB II consisting of 1M Tris-HCl (pH 7.4), 5M NaCl and 1M MgCl_2_ in sterile H_2_O containing 0.1% Tween-20 only was added to each sample on ice and the tube was inverted 3 times. Nuclei were pelleted by centrifugation at 500g for 10 min at 4°C. Supernatant was discarded carefully and pellet was resuspended in 52 µl transposition mix consisting of 25 µl 2x TD buffer, 2.5 µl transposase, 16.5 µl DPBS (+Mg, +Ca), 0.5 µl 1% Digitonin, 0.5 µl 10% Tween-20 and 5 µl H_2_O by pipetting up and down 6 times. Reaction was incubated in a thermomixer at 1000 rpm for 30 min at 37 °C and was subsequently cleaned up by using the DNA Clean and Concentrator-5 Kit (Zymo Research). For that purpose, 250 µl DNA Binding Buffer provided from the kit was added to each 50 µl sample and either stored at -20°C or further processed. Sample warmed to RT was added to the column and was centrifuged at 10000 g for 30 sec. After washing with 200 µl Wash Buffer, sample was centrifuged again at 10000 g for 30 sec and washing and centrifugation was repeated once. Sample was eluted in 22 µl Elution Buffer by centrifugation at 10000 g for 30 sec. Eluted DNA was either stored at -20 °C t or amplified by mixing 20 µl transposed sample with 25 µl 2x NEBNext Master Mix, 2.5 µl 25 µM Primer Forward (Ad1) and 2.5 µl 25 µM Primer Reverse (Ad2.1, Ad2.3-Ad2.24) and following the protocol 72 °C for 5 min, 98 °C for 30 s, 6x [98 °C for 10 s, 63 °C for 30 s, 72 °C for 1 min]. Amplified samples were purified by double sided AMPure Beads clean up. To that end, 2x concentrated Agencourt AMPure XP beads were mixed and warmed to RT for at least 30 min. 2x concentrated beads were mixed again and 27.5 µl (0.55x) was added to 50 µl PCR reaction. After mixing by pipetting up and down 10 times, sample was incubated for 10 min at RT and was subsequently placed on a magnet for 5 min to separate the beads from the solution. Cleared solution was transferred into a new tube and 17.5 µl (0.35x) 2x concentrated beads were added and mixed by pipetting up and down 10 min. After incubating 10 min at RT, sample was placed again on a magnet for 5 min. Cleared solution was discarded this time and pellet was washed with 200 µl 80% ethanol for 1 min at RT. Ethanol was discarded and washing was repeated once. After removing ethanol completely, sample was dried for 5-10 min at RT while being placed on the magnet. Off the magnet, pellet was resuspended in 20 µl low TE Buffer by pipetting up and down 10 times and incubated for 5 min at RT. After incubation, reaction was placed on the magnet for 5 min to separate the beads from the sample. 19 µl sample was transferred into a new tube and quantified by using D1000 ScreenTape on 2200 TapeStation. For that purpose, 3 µl D1000 Sample Buffer was mixed with 1 µl sample and loaded on the TapeStation. 4 ng of each sample was pooled together with samples being labelled with different barcodes due to different reverse primers ^80^ resulting in pools containing 23 samples each. Each pool was quantified again on a DNA high sensitivity bioanalyzer chip to confirm properly tagmented libraries showing a nucleosome-like banding pattern and a minimum concentration of 2 ng/µl. Lower concentrated pools were cleaned up by using the double sided AMPure Beads clean up as described above and were eluted in 17 µl low TE Buffer. Quantification on DNA high sensitivity bioanalyzer chip was repeated and 15 µl library was sent for ATAC sequencing.

#### ATACseq analysis

Paired-end raw ATAC-Seq reads were aligned against the GRCh38 genome using bowtie 2 ^81^ (version 2.3.5). Subsequently, duplicates were removed using picard tools and peak calling was performed using macs2 and standard human genome size (hs parameter). Resulting peak files of all samples were then merged using diffbind, only retaining those peaks that were present in at least 6 samples, giving rise to the union peak set employed in all further analyses. Next, bedtools multicov function was used to count fragments overlapping with peaks by at least one base. The resulting matrix peak x fragment count matrix was then further analyzed with DESeq2, using the effective library size (counting only reads overlapping with at least one peak of the union peak set) for sequencing depth normalization/size factor estimation. Subsequently, dispersions were estimate using DESeq2’s function: estimateDispersions(obj,fitType=’local’). Differential peak enrichment analysis was then performed again using a paired design in DESeq2 with the design formula ∼sex+batch+Factor and the nbinomWaldTest function. No peaks were detected differentially open between diagnosis groups below an adjusted p-value < 0.1.

### Identity verification of sequencing libraries

In order to confirm donor identity of each RNA-Seq, ATAC-Seq and microRNA-Seq library, we utilized the software verifyBam^82^, providing it with the raw bam file and a vcf file containing the genotypes of all donors. For microRNA-Seq, fastq files were re-aligned to the whole genome using STAR^63^. Identified swaps were corrected and correct labels were used in subsequent analyses. For scRNA-Seq data of iNs, we used demuxlet^73^ in combination with the aforementioned vcf file to assign individual cells to one of the four donors that were included in this experiment.

### qPCR

Realt-time PCR was performed for experiments in manuscript. For a detailed primer list see key resources table.

#### Sample collection

Samples were prepared as described above (4.1.1) and further processed for cDNA synthesis by using RevertAid H Minus First Strand cDNA Synthesis Kit (Thermo Fisher Scientific).

#### cDNA synthesis

Oligo(dT)18 primer (1µl from 100 µM stock) was added to 500 ng RNA filled up to 11 µl with DNA/RNAse-free H_2_O. After incubation for 5 min at 65 °C, the reaction was kept on ice. 4 µl 5x Reaction Buffer, 1 µl RiboLock RNase Inhibitor (20 U/ µl), 2 µl 10 mM dNTP Mix and 1 µl RevertAid H Minus Reverse Transcriptase (200U/ µl) was added to each reaction, incubated for 1 hr at 42 °C and stored at -20 °C or directly used for qPCR. For that purpose, 5 µl 2x SSO Advanced Biorad Mastermix, 0.25 µl 10 µM Primer Forward, 0.25 µl 10 µM Primer Reverse and 3.5 µl H_2_O were mixed with 1 µl cDNA diluted 1:10 in H_2_O and loaded on LightCycler 480 II from Roche or BIO-RAD CFX thermal cycler following the protocol 95 °C for 2 min, 35x [95 °C for 10 s, 62 °C for 15 s, 72 °C for 5 s], 60°C for 1 s (2.5 °C/ s), 95 °C continuous (0,11 °C/ s), 45 °C for 1 s.

#### Analysis

For qPCR analysis, CT-values of target gene and reference (e.g. MAP2, RTF2 or Distal/Proximal UTR, Supplementary Table 4) were used to calculate target specific relative expression as 2^ (-(CT_target_-CT_Reference_)) giving rise to normalized transcript levels.

The distal/proximal 3’UTR fold change for the dAPA locus specific analysis was calculated as the arithmetic ratio between the relative expression of the two different 3’UTR regions belonging to the same gene locus.

The normalized transcript levels were analyzed by linear mixed models as implemented in the lme4 R package^83^ modeling diagnosis as fixed effect and batch as well as donor (for APA comparison) as random effect. Significance of the fixed effect was assessed using the two-tailed t-test as implemented lmerTest package relying Satterthwaite’s method for approximating degrees of freedom^84^. qPCR value distribution as visualized as violin or boxplot. For visualization purposes, we normalized transcript values for each batch by dividing the respective transcript abundance levels by the mean across all control samples from entire batch plotted the results.

### Western blot

#### Harvesting

iNs were collected at D 49 in 1.4 ml cold DPBS in a 1.5 ml reaction tube and centrifuged at 300 g for 5 min. Supernatant was discarded and cell pellet was stored at -80 °C until further use.

#### Protein extraction

Cell pellets were thawed on ice and centrifuged for 5 min at 500 x g and 4°C. Remaining DPBS supernatant was removed by pipetting. Then pellets were washed once with ice-cold DPBS. After complete aspiration of the supernatant (DPBS), ice-cold lysis buffer/RIPA buffer containing 150 mM NaCl, 1% NP-40, 0.1% SDS, 50 mM Tris Ph8.1, 0.5% sodium deoxycholate and proteinase inhibitor was added and incubated 5 min on ice. After homogenization by pipetting up and down 5x, samples were incubated additional 30 min with brief vortexing every 5 min. Lysated samples were then centrifuged for 15 min at 14,000 x g and 4°C. Immediately after centrifugation, supernatant was transferred to a clean microfuge tube and the concetration was analyzed by Bradford assay (Biorad) using a BSA standard-curve as reference. Protein extract concentrations were uniformed to 1µg/µl in lysis buffer and 5µg aliquots were stored at -80°C.

#### Digital western blotting

Sample preparation for digital western blotting using Wes™ (ProteinSimple, USA, PSD95 experiments) or Abby (ProteinSimple, USA, PTBP2 KD experiments) was performed according to manufactures instructions. First, 400mM DTT solution from EZ Standard Pack 1 was prepared by addition of 40µl H20 and Fluorescent 5X Master Mix was prepared by addition of 20µl 10X Sample Buffer and 20µl 400mM DTT solution. Primary antibody dilutions were prepared using Antibody Diluent 2 (PSD95 1:200, b3TUB 1:50). For each sample, one part (1µl) of 5X Fluorescent Master Mix and four parts of protein lysates (adjusted with 0.1X Sample Buffer to a total protein amount of: b3TUB, 0.1µg; PSD95, 1.6µg) were gently mixed, vortexed, denaturated at 95°C for 5 min, vortexed, spined down, and stored on ice before loading. Pre-filled Plates (12-230 kDa) were loaded with Antibody Diluent 2, Streptavidin-HRP, Anti-Mouse Secondary HRP Antibody or Anti-Rabbit Secondary HRP Antibody according to primary antibody application, Biotinylated Ladder, freshly prepared mix of Luminol-S and Peroxide, and Wash Buffer according to manufacturer’s instructions. Plates were centrifuged for 5 min at 1000g at RT. Digital western blotting with prepared 12-230 kDa plates and 25-Capillary cartridges for Size based Separation at Wes™ was performed with default settings.Subsequent analysis were performed with the “Compass” software (Compass for SW 5.0.1 Mac Beta), on the best exposure time for each individual run, selected based on to chemiluminescence intensity linearity. determination of the peak area was performed using ‘Dropped line’. Data were exported as Excel files for subsequent statistical analysis.

#### Analysis

PSD95 intensity values for the high molecular weight (90-95 kDa) were divided by b3Tub intensity from the same protein extract and log_2_ transformed, giving rise to the normalized intensity ratio (NRI). For PTBP2, intensity values were divided by the intensity of histone 3 (H3) from the same nuclear extracts. Statistical testing of NRIs (**Supplementary Table 4**) was conducted using by linear mixed models as implemented in the lme4 R package^83^ modeling diagnosis or treatment as fixed effect and batch as well as donor (for PSD95 analysis) as random effect. Significance of the fixed effect was assessed using the t-test as implemented lmerTest package relying Satterthwaite’s method for approximating degrees of freedom^84^. Prior to analysis, outliers were filtered using Tukey’s outlier test (including only values within 1.5 x inter-quartile range) in a batch specific manner for each variable separately.

For visualization purposes, we normalized the NRI values for each batch by dividing the respective NRI values by the mean NRI from control samples across the entire batch and plotted the results.

### Immunocytochemistry (ICC)

In order to investigate differential gene expression on protein level regarding quantity and localization, immunofluorescent stainings were performed.

#### Fixation and staining

Samples were washed twice with DPBS, fixed with 4% Paraformaldehyde for 15 min at RT and washed again twice with DPBS. After permeabilization with 0.1% Triton-X-100 in DPBS for 15 min at RT, cells were blocked with 1% BSA and 1% donkey serum in 0.1% Triton-X-100 for 45 min at RT. Primary antibody was added in blocking solution over night at 4 °C. Cells were washed three times with 0.1% Triton-X-100, and incubated with secondary antibody in blocking solution for 2.5 hr in the dark at RT. Cells were washed twice with 0.1 Triton-X-100, once with DPBS and mounted in Aqua-Poly/Mount from Polysciences™.

#### iNs maturity analysis

iNs from Ctrls (n=10) and ISCZ (n=8) were fixed and stained with DAPI, FOXG1, and CUX1 as described above. Primary image analysis and segmentation was performed using CellInsight CX5 High-Content Screening System and HCS Navigator^TM^ Version 6.6.1 Software from Thermo Scientific^TM^. Briefly, 3 wells per donor (2 donors with 2 wells, 2 donors with 1 well per donor) were investigated and 9 fields of view (FOV) were taken at randomly defined but fixed positions for each well at 20X magnification. Subsequently each image was segmented for the imaging analysis using Fiji^59^ and the BIOVOXXEL toolbox^85^, see also https://github.com/biovoxxel, based on ImageJ, Version 1.15f51. First, nuclei, FOXG1 and CUX1 segmentation masks were separately applied using the appropriate fluorescent channels. Then, expression of FOXG1 or CUX1 within a certain cell was defined by an overlap of the FOXG1 or CUX1-segmentation mask with the DAPI-based nuclei segmentation-defined regions of interests. Number of total cells (DAPI-based segmentation) and number of cells that express FOXG1 (overlap DAPI segmentation and FOXG1 segmentation) and CUX1 (overlap DAPI segmentation and CUX1 segmentation) were exported per field of view. Results are shown in **Extended Data Fig. 2b,c**.

#### Synapse count analysis

To measure synapse number in iNs from Ctrls (n=10) and ISCZ (n=10), iNs were fixed and stained as described above. Synapses were defined by pixel overlaps of the synaptic markers Synapsin (SYN1) and the neurite/neuronal marker βTubulin. Primary image analysis and segmentation was performed using CellInsight CX5 High-Content Screening System and HCS Navigator^TM^ Version 6.6.1 Software from Thermo Scientific. Briefly, 30 fields of view (FOV) were taken at randomly defined but fixed positions for each well at 20X magnification. Subsequently, each image was segmented into cell body and neurite compartments using the DAPI and βTubulin channel in combination with morphological and intensity parameters such as size, shape and differences in pixel intensities.

For that purpose, nuclei were recognized in the DAPI channel while cell bodies were identified by βTubulin staining overlapping with tagged nuclei at a minimum of 10 %. Neurite segmentation mask was generated by using βTubulin channel as well. SYN1 spots were identified by aforementioned parameter settings and one additional channel showing SYN1 staining was determined. Overlap of puncta was defined as a minimal intersection of 10 % within the cell body or neurite segmentation masks defined by βTubulin staining^86^.

The detailed protocol for segmentation and analysis of CellInsight software is available upon request as XML file.

Following primary feature segmentation, definition of relevant SYN1 puncta was performed by intersecting the latter feature masks with those of detected cell bodies and neurites. Only puncta overlapping with the latter were considered for further analysis. Next, measurements were collapsed for each FOV, computing the SYN1 puncta density in neurites and cell bodies separately, by dividing the total number or area of the former puncta per FOV by the total area of either cell bodies or neurites. Prior to further analysis, FOV were filtered for a minimal and maximal detected cell body and neurite area, excluding FOVs at the extremes of the distribution (excluding empty FOVs and high cell body density FOVs). Finally, the resulting density values per FOV were averaged for each well and used for subsequent statistical analysis.

For latter, we used linear mixed models as implemented in the lme4 R package^83^ modeling diagnosis as fixed effect and batch as well as donor as random effect. Significance of the fixed effect was assessed using the t-test as implemented lmerTest package relying Satterthwaite’s method for approximating degrees of freedom^84^. Model fit, residual distribution and diagnostics were evaluated using the R package DHARMa^87^. All well level density values used can be found in **Supplementary Table 4**.

### PTBP2 knockdown

To functionally assess the contribution of PTBP2 to the dAPA in iNs from ISCZ, PTBP2 repression was modulated using shRNAs. To that end, lentiviral particles containing 4 different shRNAs (**key resources table**) targeting the human PTBP2 transcript (TRC database) or a non-targeting control lentivirus were produce using the protocol described above. In order to limit the effect the effects of PTBP2 knockdown on neuronal development/maturation and rather assess its relevance for o neuronal physiology under steady state conditions, we performed an acute knockdown towards the end of the differentiation timeline. We evaluated distinct transduction time-points and duration between day 30 and 65 testing 5, 10 and 15 days of culture under knockdown conditions to exclude any dependency of the silencing effect on the differentiation stage, collecting iNs at day 44, 49, 62 and 70. Evaluation of PTBP2 knockdown magnitude and impact on PSD95 protein levels showed no dependency on transduction time point or duration and data points for PTBP2 knockdown or PSD95 were pooled into a Ctrl and PTBP2 knockdown condition. Based on these observations, we selected 15 days of knockdown duration and collection at day 49 for synapse density analysis, considering the possibility of a lag in PTBP2 knockdown effect on synapse turnover. Accordingly, iNs were infected in BrainPhys media supplemented with 6µg/ml Polybrene and cultured following the regular protocol.

#### Synaptic density analysis in PTBP2^KD^

To determine synaptic density changes induced by the downregulation of PTBP2, iNs were fixed with 4% PFA in DPBS for 10 min at RT, permeabilized with 0.2% Triton-X in DPBS for 10 min and unspecific epitope recognition was blocked using 0.5% Tween-20 and 1% BSA in DPBS for 60 min. Primary antibody (anti-Syn1 and anti-Tubb3) were incubated in the blocking solution supplemented with 1% normal donkey serum overnight at 4°C, the excess of antibody was washed 3 times with 0.5% Tween-20 in DPBS for 5 min each wash. Secondary antibodies were incubated in blocking solution supplemented with 1% normal donkey serum for 60 min at room temperature. Stained cells were maintained in Aqua-Poly/Mount media at 4°C.

Imaging acquisition was automatized on the CellInsight CX5 High-Content Screening system using 20x magnification objective (20 field/well, 4 well/condition) with 2048dpi. TIFF file were exported and analyzed using CellProfiler 4.2.1. Nuclei were detected based on the DAPI channel using the Adaptive Otsu thresholding method for intensities between 0.5 and 0.9. iN nuclei were detected and distinguished from the co-cultured mouse astrocytes based on size (diameter between 24 and 70 pixels) and on the absence of chromocenter structures (FilterObjects by Texture_Correlation and by AreaShape_Compactness). The identified nuclei were then further used for the identification of the soma as secondary objects (Distance – B method, with 60 pixels as maximum distance) using the Tubb3 signal.

To avoid any exclusion of soma regions, primary object identification was also performed on the TUBB3 channel using the Adaptative Otsu thresholding method with intensities between 0.48 and 0.9 and a permissive size interval between 30 and 300 pixels. The two independently detected soma maks were then combined into a final comprehensive soma mask. Neurite tubeness was enhanced with a smoothing scale of 1 for features with 20 pixels of maximum size to facilitate the identification of a partial network (including some soma regions) using the Adaptative with Robust background thresholding method for intensities between 0.0115 and 0.04 with a typical diameter between 6 and 400 pixels. Removing from the latter mask the final soma mask we obtained the Neurite compartment mask. The enhancement of speckles with max size of 5 pixel facilitated the primary objects identification for Syn1 signals(Adaptative Otsu, intensity 0.04,0.3) Detected Synapsin objects within the different compartments (Soma and Neurites) were counted, and the occupied area of all the masks and sub-masks was calculated.

### Electrophysiology

#### Patch clamp

Whole-cell current- and voltage-clamp recordings (−70 mV holding potential, >1 GΩ seal resistance, <20 MΩ series resistance, 8 mV liquid junction potential correction, 3 kHz low-pass filter, 15 kHz sampling rate) from cultured neurons were conducted at room temperature (23-25°C) using an EPC9 amplifier (HEKA). Cells were superfused (2-3 ml/min flow rate) with a carbogen gas (95% O2/5% CO2)-saturated solution containing (in mM): 121 NaCl, 4.2 KCl, 29 NaHCO3, 0.45 NaH2PO4, 0.5 Na2HPO4, 1.1 CaCl2, 1 MgSO4, and 20 D-glucose (Bardy et al., 2015). This solution additionally contained NBQX (5 µM) and picrotoxin (100 µM) for current-clamp measurements. For current-clamp measurements, patch pipettes (3-5 MΩ open tip resistance) were filled with a solution consisting of (in mM): 135 KMeSO4, 8 NaCl, 0.3 EGTA, 10 HEPES, 2 Mg-ATP, and 0.3 Na-GTP. Current injections were used to depolarize or hyperpolarize the neuron under investigation. Offline analysis was performed using the Mini Analysis Program (version 6.0.7, Synaptosoft).

### Quantitative trait analysis

#### Expression quantitative trait (eQTL) analysis

For eQTLs analysis, RNA-Seq reads were re-aligned to hg19 using the same strategy as outlined in the section RNA-Seq processing, since all genotype-based analyses were conducted on hg19. We utilized the gtex-pipeline implemented by the Broad institute for the analysis to preprocess the data before identifying the eQTLs by linear regression with FastQTL^88^. We only included genes with ≥ 0.1 TPM and ≥ 6 unnormalized reads in ≥ 20% of the samples. Thereafter each of the remaining 24,799 genes was inverse normal transformed across samples. We calculated 15 PEER factors from the resulting expression data, which were used as the covariates, as well as the sex and differentiation site of each sample for FastQTL.

We identified cis-EQTLs within a 200 kb window of the Transcription Start Site of each included gene and by performing both a nominal and an adaptive permutation pass, running between 1000 and 10000 permutations, of FastQTL on the data. We also computed the allelic Fold Change as a comparative measurement of effect size of cis-eQTLs^89^. Subsequently, eQTLs with a multiple testing corrected permutation based p-value below 0.05 were defined as significant and considered for further analysis consistent with the GTEx approach.

#### microRNA quantitative trait (mirQTL) analysis

The mirQTL analysis was done analogous to the eQTL analysis, using the GTEx preprocessing pipeline and FastQTL. We kept genes with ≥ 0.1 TPM and ≥ 3 unnormalized reads in ≥ 10% of samples, resulting in 832 remaining mature miRNAs. The covariates consisted of the first 3 PEER factors, omitting the remaining, colinear factors, 3 genotype PCs, 3 miRNA PCs and the sex and differentiation site of each sample. We ran both a nominal and adaptive permutation pass on the data, keeping the parameters of the eQTL analyses, but setting the FDR to 0.2.

#### Chromatin quantitative trait (caQTL) analysis

For caQTLs analysis, ATAC-Seq reads were re-aligned to hg19 using the same strategy as outlined in the section ATAC-Seq processing since all genotype based analyses were conducted on hg19. Newly aligned bam files were then used for caQTL mapping in combination with RASQUAL^90^. To that end, we created a fragment count table based on the previously identified union peak set lifted over to hg19 using the UCSC liftover tool (http://genome.ucsc.edu/cgi-bin/hgLiftOver)^91^ and bedtools multicov (see ATAC-Seq data processing). Subsequently, we created a modified VCF file containing the allele-specific counts derived from the master VCF file using the RASQUAL toolbox script createASVCF.sh. We only included variants in a 200 kb *cis* window of the transcription starting sites of each peak and added the -z flag to convert the genome imputation quality score to allelic probability and the –force flag to include fitting the model for large numbers of fSNPs and rSNPs, accepting the added computational effort. As covariate, we included sex, differentiation site and the first PC of the genotype PCA. In addition, we ran RASQUAL with the-r flag to generate a random permutation of each feature and obtain empirical null distribution of p-values. Both the original and permuted output values were further processed using eigenMT^92^ to adjust for multiple testing and compare the experimental p-values to the null distribution. To that end, RASQUAL based p-values were Bonferroni corrected for each peak, using the number of SNPs tested for each peak. Subsequently, different Bonferroni corrected p-value thresholds were used to assess the number of peas with at least one significant caQTL in the original and permuted dataset. Based on this analysis, we selected a Bonferroni corrected p-value threshold of 0.01 yielding an empirical FDR of 0.16 to define the set of significant caQTL peaks.

### Overlap of QTLs with human post mortem and GWAS data

For overlap analysis with GWAS data, we downloaded the SCZ GWAS summary statistics from Pardinas et al.^33^ from the PGC website and intersected genomic coordinates of eQTLs/caQTLs/mirQTLs on hg19. The results are shown in **Fig. 1**. For intersection with eQTLs from post mortem brain material, we downloaded eQTL summary results^93^ from the common mind consortium^11^ (https://www.synapse.org/#!Synapse:syn2759792/wiki/69613) and intersected the genomic coordinates.

### RBP binding motif analysis in 3’UTR sequences

For RBP motif enrichment with the UTR of transcripts subject to APA, we performed computational motif enrichment using *Transite*^94^. The latter that leverages a large collection of available RBP and expression data from the ENCODE project. We performed *Transite* analysis for transcript with increased or decreased distal APA site usage in SCZ separately, using the entire set of 3’UTR from all gencode genes as background. Resulting enrichment scores and p-values for the set of transcripts with increased usage of the distal PAS are depicted in Fig. 5c with p-values corrected for multiple testing using the q-value method and listed in Supplementary Table 3. Enrichment analysis for RBP binding site in the set of transcripts with more proximal UTR usage in SCZ yielded no significant results.

## Data availability

The data for this project have been deposited in GEO under accession [TBA] or the European genome phenome archive, accession number [TBA] for privacy protected data.

## Materials & Correspondence

Materials used in this manuscript are available through Michael J. Ziller and Moritz J. Rossner, conditional on proper ethics approval from the requesting institution. A subset of iPSC lines is subject to sharing constraints due to limited donor consent.

## Acknowledgments

First, we would like to thank the patients for their study participation and tissue donation. We would like to thank the entire Ziller and Rossner lab team as well as many colleagues from the Department of Psychiatry at the LMU Munich, Germany, and the Department of Translational Research in Psychiatry, MPIP, Germany. We acknowledge the expert help of the Douglas–Bell Canada Brain Bank staff (J. Prud’homme, M. Bouchouka and A. Baccichet), and H. Djambazian at the MUGQIC. Moreover, we thank Jan Brocher from BioVoxxel for his excellent help with imaging analysis and Lucio Di Filippo for his support of the microRNA library preparation.

This study used data from the CommonMind consortium provided through NIMH. Data for this publication were obtained from NIMH Repository & Genomics Resource, a centralized national biorepository for genetic studies of psychiatric disorders. Data were generated as part of the CommonMind Consortium supported by funding from Takeda Pharmaceuticals Company Limited, F. Hoffman-La Roche Ltd and NIH grants R01MH085542, R01MH093725, P50MH066392, P50MH080405, R01MH097276, RO1-MH-075916, P50M096891, P50MH084053S1, R37MH057881, AG02219, AG05138, MH06692, R01MH110921, R01MH109677, R01MH109897, U01MH103392, and contract HHSN271201300031C through IRP NIMH. Brain tissue for the study was obtained from the following brain bank collections: the Mount Sinai NIH Brain and Tissue Repository, the University of Pennsylvania Alzheimer’s Disease Core Center, the University of Pittsburgh NeuroBioBank and Brain and Tissue Repositories, and the NIMH Human Brain Collection Core. CMC Leadership: Panos Roussos, Joseph Buxbaum, Andrew Chess, Schahram Akbarian, Vahram Haroutunian (Icahn School of Medicine at Mount Sinai), Bernie Devlin, David Lewis (University of Pittsburgh), Raquel Gur, Chang-Gyu Hahn (University of Pennsylvania), Enrico Domenici (University of Trento), Mette A. Peters, Solveig Sieberts (Sage Bionetworks), Thomas Lehner, Stefano Marenco, Barbara K. Lipska (NIMH).

This work was supported by BMBF, eMed grant numbers 01ZX1504, 01ZX1706A (MJZ), Else-Kroener-Fresenius Stiftung, grant A54 (MJZ), Research grants provided by Boehringer Ingelheim and the BMBF (FKZ 1EK2101D) (MJR), the Näder Foundation (PF, MJR), Else Kröner-Fresenius Research School for Translational Psychiatry (FJR), Munich Clinician Scientist Program (MCSP), FöFoLe 009/2019 (FJR), German Research Foundation, grants FOR2107 DA1151/5-1 and DA1151/5-2 (UD), Intramural Research Program of the NIMH, ZIA-MH002810 (FC), Italian Ministry of Health (Piano Operativo Salute Traiettoria 3, “Genomed”), Fondazione Telethon Core Grant, Armenise-Harvard Foundation Career Development Award, European Research Council (grant agreement 759154, CellKarma), and the Rita-Levi Montalcini program from MIUR (DC), and Rita-Levi Montalcini program from MIUR (MC).

## Author contributions

iPSC generation: AH, FJR., AH, MG, CR, SG, NG, SM, VS, VA; iPSC characterization: AH, FJR, RA, AH, CR, SM, LT, LJB, MRH, SP, SG, NG, VA, SS, MW; Differentiation experiments: AH, FJR, SG, NG, AH, BK; Omics profiling: AH, MG, SR, RA, AH, CR, MR, DC; Molecular biology experiments: AH, FJR, MG; Imaging and imaging analysis: AH, FJR, MG, RA, SM, AA, MJZ; Electrophysiology: AH, DM, BH, EMW, VS, ME; Bioinformatics: LT, LW, VM, LJB, MJZ; Planned experiments: AH, FJR, MG; Fibroblast collection/processing: CR, AH; Provided critical resources incl. PBMC collection and patient characterization: FR, SG, NG, IP, TOS, SDW, FJM, AS, PF, AH, UD, IN, TK, GT, EBB; Data visualization and curation: FJR, AH and MJZ with the help of all authors; Wrote the paper: MJZ, FJR, DS, AH with the help of all authors; Critically revised the manuscript: EBB, MJR; Acquired Funding: MJZ, MJR; Supervised the work: MJZ, MJR; Conceived study, designed experiments and analyses: MJZ.

## Competing interests

SG and MCW are part-time employees by and shareholders of Systasy Bioscience GmbH, Munich, Germany. MJR is shareholder and consultant of Systasy Bioscence GmbH. DC is founder, shareholder, and consultant of NEGEDIA (Next Generation Diagnostic) is founder, shareholder, and consultant of Next Generation Diagnostic srl. Sara Riccardo is an employee of NEGEDIA. All other authors declare that they have no competing interests.

## Notes

### Summary of Updates

Minor: Corrected gene name spelling errors

## References

1. Charlson, F.J. et al. Global Epidemiology and Burden of Schizophrenia: Findings From the Global Burden of Disease Study 2016. Schizophr Bull 44, 1195–1203 (2018).

2. Hilker, R. et al. Heritability of Schizophrenia and Schizophrenia Spectrum Based on the Nationwide Danish Twin Register. Biol Psychiatry 83, 492–498 (2018).

3. Trubetskoy, V. et al. Mapping genomic loci implicates genes and synaptic biology in schizophrenia. Nature 604, 502–508 (2022).

4. Singh, T. et al. Rare coding variants in ten genes confer substantial risk for schizophrenia. Nature 604, 509–516 (2022).

5. Howard, D.M. et al. Genome-wide meta-analysis of depression identifies 102 independent variants and highlights the importance of the prefrontal brain regions. Nat Neurosci 22, 343–352 (2019).

6. Mullins, N. et al. Genome-wide association study of more than 40,000 bipolar disorder cases provides new insights into the underlying biology. Nat Genet 53, 817–829 (2021).

7. The Network and Pathway Analysis Subgroup of the Psychiatric Genomics Consortium. Psychiatric genome-wide association study analyses implicate neuronal, immune and histone pathways. Nat Neurosci 18, 199–209 (2015).

8. Wang, D. et al. Comprehensive functional genomic resource and integrative model for the human brain. *Science (New York*, N.Y.) 362, eaat8464 (2018).

9. Schrode, N. et al. Synergistic effects of common schizophrenia risk variants. Nat Genet 51, 1475–1485 (2019).

10. Sekar, A. et al. Schizophrenia risk from complex variation of complement component 4. Nature 530, 177–83 (2016).

11. Fromer, M. et al. Gene expression elucidates functional impact of polygenic risk for schizophrenia. Nat Neurosci 19, 1442–1453 (2016).

12. Bryois, J. et al. Evaluation of chromatin accessibility in prefrontal cortex of individuals with schizophrenia. Nat Commun 9, 3121 (2018).

13. Gandal, M.J. et al. Transcriptome-wide isoform-level dysregulation in ASD, schizophrenia, and bipolar disorder. Science 362(2018).

14. Dobbyn, A. et al. Landscape of Conditional eQTL in Dorsolateral Prefrontal Cortex and Co-localization with Schizophrenia GWAS. Am J Hum Genet 102, 1169–1184 (2018).

15. Mayr, C. What Are 3’ UTRs Doing? Cold Spring Harb Perspect Biol 11(2019).

16. Li, L. et al. An atlas of alternative polyadenylation quantitative trait loci contributing to complex trait and disease heritability. Nat Genet 53, 994–1005 (2021).

17. Park, C.Y. et al. Genome-wide landscape of RNA-binding protein target site dysregulation reveals a major impact on psychiatric disorder risk. Nat Genet 53, 166–173 (2021).

18. Kircher, T. et al. Neurobiology of the major psychoses: a translational perspective on brain structure and function-the FOR2107 consortium. Eur Arch Psychiatry Clin Neurosci 269, 949–962 (2019).

19. Bruckl, T.M. et al. The biological classification of mental disorders (BeCOME) study: a protocol for an observational deep-phenotyping study for the identification of biological subtypes. BMC Psychiatry 20, 213 (2020).

20. Krcmar, L. et al. The multimodal Munich Clinical Deep Phenotyping study to bridge the translational gap in severe mental illness treatment research. Front Psychiatry 14, 1179811 (2023).

21. Budde, M. et al. A longitudinal approach to biological psychiatric research: The PsyCourse study. Am J Med Genet B Neuropsychiatr Genet (2018).

22. Lee, P.H. et al. Genomic Relationships, Novel Loci, and Pleiotropic Mechanisms across Eight Psychiatric Disorders. Cell 179, 1469–1482.e11 (2019).

23. Anttila, V. et al. Analysis of shared heritability in common disorders of the brain. Science 360(2018).

24. Hoffmann, A., Ziller, M. & Spengler, D. Childhood-Onset Schizophrenia: Insights from Induced Pluripotent Stem Cells. Int J Mol Sci 19(2018).

25. Brennand, K.J. et al. Modelling schizophrenia using human induced pluripotent stem cells. Nature 473, 221–5 (2011).

26. Nehme, R. et al. Combining NGN2 Programming with Developmental Patterning Generates Human Excitatory Neurons with NMDAR-Mediated Synaptic Transmission. Cell Rep 23, 2509–2523 (2018).

27. Baron, M. et al. A Single-Cell Transcriptomic Map of the Human and Mouse Pancreas Reveals Inter- and Intra-cell Population Structure. Cell Syst 3, 346–360 e4 (2016).

28. Howard, D. et al. An in vitro whole-cell electrophysiology dataset of human cortical neurons. Gigascience 11(2022).

29. Yuan, W. et al. Temporally divergent regulatory mechanisms govern neuronal diversification and maturation in the mouse and marmoset neocortex. Nat Neurosci 25, 1049–1058 (2022).

30. Geaghan, M. & Cairns, M.J. MicroRNA and Posttranscriptional Dysregulation in Psychiatry. Biol Psychiatry 78, 231–9 (2015).

31. Forrest, M.P. et al. Open Chromatin Profiling in hiPSC-Derived Neurons Prioritizes Functional Noncoding Psychiatric Risk Variants and Highlights Neurodevelopmental Loci. Cell Stem Cell 21, 305–318 e8 (2017).

32. Flaherty, E. et al. Neuronal impact of patient-specific aberrant NRXN1alpha splicing. Nat Genet 51, 1679–1690 (2019).

33. Pardinas, A.F. et al. Common schizophrenia alleles are enriched in mutation-intolerant genes and in regions under strong background selection. Nat Genet 50, 381–389 (2018).

34. Hauberg, M.E., Roussos, P., Grove, J., Borglum, A.D., Mattheisen, M. & Schizophrenia Working Group of the Psychiatric Genomics, C. Analyzing the Role of MicroRNAs in Schizophrenia in the Context of Common Genetic Risk Variants. JAMA Psychiatry 73, 369–77 (2016).

35. Yao, Y. et al. Cell type-specific and cross-population polygenic risk score analyses of MIR137 gene pathway in schizophrenia. iScience 24, 102785 (2021).

36. You, X. et al. Investigating aberrantly expressed microRNAs in peripheral blood mononuclear cells from patients with treatmentresistant schizophrenia using miRNA sequencing and integrated bioinformatics. Mol Med Rep 22, 4340–4350 (2020).

37. Yu, H.-c., et al. Alterations of miR-132 are novel diagnostic biomarkers in peripheral blood of schizophrenia patients. Progress in Neuro-Psychopharmacology and Biological Psychiatry 63, 23–29 (2015).

38. Nanou, E. & Catterall, W.A. Calcium Channels, Synaptic Plasticity, and Neuropsychiatric Disease. Neuron 98, 466–481 (2018).

39. Tian, B. & Manley, J.L. Alternative polyadenylation of mRNA precursors. Nat Rev Mol Cell Biol 18, 18–30 (2017).

40. Shah, A., Mittleman, B.E., Gilad, Y. & Li, Y.I. Benchmarking sequencing methods and tools that facilitate the study of alternative polyadenylation. Genome Biol 22, 291 (2021).

41. Lopez-Murcia, F.J., Reim, K., Jahn, O., Taschenberger, H. & Brose, N. Acute Complexin Knockout Abates Spontaneous and Evoked Transmitter Release. Cell Rep 26, 2521–2530 e5 (2019).

42. Mitok, K.A., Keller, M.P. & Attie, A.D. Sorting through the extensive and confusing roles of sortilin in metabolic disease. J Lipid Res 63, 100243 (2022).

43. Finucane, H.K. et al. Heritability enrichment of specifically expressed genes identifies disease-relevant tissues and cell types. Nat Genet 50, 621–629 (2018).

44. Hwang, H.W. et al. cTag-PAPERCLIP Reveals Alternative Polyadenylation Promotes Cell-Type Specific Protein Diversity and Shifts Araf Isoforms with Microglia Activation. Neuron 95, 1334–1349 e5 (2017).

45. Zheng, S., Gray, E.E., Chawla, G., Porse, B.T., O’Dell, T.J. & Black, D.L. PSD-95 is post-transcriptionally repressed during early neural development by PTBP1 and PTBP2. Nat Neurosci 15, 381–8, S1 (2012).

46. Koopmans, F. et al. SynGO: An Evidence-Based, Expert-Curated Knowledge Base for the Synapse. Neuron 103, 217–234 e4 (2019).

## Methods-only references

15. Mayr, C. What Are 3’ UTRs Doing? Cold Spring Harb Perspect Biol 11(2019).

34. Hauberg, M.E. et al. Analyzing the Role of MicroRNAs in Schizophrenia in the Context of Common Genetic Risk Variants. JAMA Psychiatry 73, 369–77 (2016).

45. Beveridge, N.J. & Cairns, M.J. MicroRNA dysregulation in schizophrenia. Neurobiol Dis 46, 263–71 (2012).

47. Okita, K. et al. An efficient nonviral method to generate integration-free human-induced pluripotent stem cells from cord blood and peripheral blood cells. Stem Cells 31, 458–66 (2013).

48. Chou, B.K. et al. A facile method to establish human induced pluripotent stem cells from adult blood cells under feeder-free and xeno-free culture conditions: a clinically compliant approach. Stem Cells Transl Med 4, 320–32 (2015).

49. Diecke, S. et al. Novel codon-optimized mini-intronic plasmid for efficient, inexpensive, and xeno-free induction of pluripotency. Sci Rep 5, 8081 (2015).

50. Danecek, P., McCarthy, S.A., HipSci, C. & Durbin, R. A Method for Checking Genomic Integrity in Cultured Cell Lines from SNP Genotyping Data. PLoS One 11, e0155014 (2016).

51. Danecek, P. et al. Twelve years of SAMtools and BCFtools. Gigascience 10(2021).

52. Guo, Y. et al. Illumina human exome genotyping array clustering and quality control. Nat Protoc 9, 2643–62 (2014).

53. Kilpinen, H. et al. Common genetic variation drives molecular heterogeneity in human iPSCs. Nature 546, 370–375 (2017).

54. Wray, N.R. et al. Genome-wide association analyses identify 44 risk variants and refine the genetic architecture of major depression. Nat Genet 50, 668–681 (2018).

55. Ge, T., Chen, C.Y., Ni, Y., Feng, Y.A. & Smoller, J.W. Polygenic prediction via Bayesian regression and continuous shrinkage priors. Nat Commun 10, 1776 (2019).

56. Zarrei, M. et al. A large data resource of genomic copy number variation across neurodevelopmental disorders. NPJ Genom Med 4, 26 (2019).

57. Marshall, C.R. et al. Contribution of copy number variants to schizophrenia from a genome-wide study of 41,321 subjects. Nat Genet 49, 27–35 (2017).

58. Kent, W.J. et al. The human genome browser at UCSC. Genome Res 12, 996–1006 (2002).

59. Schindelin, J., et al. Fiji: an open-source platform for biological-image analysis. Nat Methods 9, 676–82 (2012).

60. Bock, C. et al. Reference Maps of human ES and iPS cell variation enable high-throughput characterization of pluripotent cell lines. Cell 144, 439–52 (2011).

61. Tsankov, A.M. et al. A qPCR ScoreCard quantifies the differentiation potential of human pluripotent stem cells. Nat Biotechnol 33, 1182–92 (2015).

62. Zhang, Y. et al. Rapid single-step induction of functional neurons from human pluripotent stem cells. Neuron 78, 785–98 (2013).

63. Dobin, A. et al. STAR: ultrafast universal RNA-seq aligner. Bioinformatics 29, 15–21 (2013).

64. Rummel, C.K. et al. Cell type and condition specific functional annotation of schizophrenia associated non- coding genetic variants. bioRxiv (2023).

65. Karapetyan, A. transcriptR: An Integrative Tool for ChIP- And RNA-Seq Based Primary Transcripts Detection and Quantification. (2022).

66. Szkop, K.J., Moss, D.S. & Nobeli, I. flexiMAP: a regression-based method for discovering differential alternative polyadenylation events in standard RNA-seq data. Bioinformatics 37, 1461–1464 (2021).

67. Xia, Z. et al. Dynamic analyses of alternative polyadenylation from RNA-seq reveal a 3’-UTR landscape across seven tumour types. Nat Commun 5, 5274 (2014).

68. Storey, J.D. The positive false discovery rate: A Bayesian interpretation and the q-value. Annals of Statistics 31, 2013–2035 (2003).

69. Quinlan, A.R. & Hall, I.M. BEDTools: a flexible suite of utilities for comparing genomic features. Bioinformatics 26, 841–2 (2010).

70. Bulik-Sullivan, B. et al. An atlas of genetic correlations across human diseases and traits. Nat Genet 47, 1236–41 (2015).

71. Xie, Z. et al. Gene Set Knowledge Discovery with Enrichr. Curr Protoc 1, e90 (2021).

72. Zhao, S., Li, C.I., Guo, Y., Sheng, Q. & Shyr, Y. RnaSeqSampleSize: real data based sample size estimation for RNA sequencing. BMC Bioinformatics 19, 191 (2018).

73. Kang, H.M. et al. Multiplexed droplet single-cell RNA-sequencing using natural genetic variation. Nat Biotechnol 36, 89–94 (2018).

74. Team, R.C. R: A Language and Environment for Statistical Computing. (R Foundation for Statistical Computing, Vienna, Austria, 2019).

75. Stuart, T. et al. Comprehensive Integration of Single-Cell Data. Cell 177, 1888–1902 e21 (2019).

76. Kryuchkova-Mostacci, N. & Robinson-Rechavi, M. A benchmark of gene expression tissue-specificity metrics. Brief Bioinform 18, 205–214 (2017).

77. Krueger, F.J.F.; Ewels, P.; Afyounian, E.; Schuster-Boeckler, B. TrimGalore. (2021).

78. Langmead, B., Trapnell, C., Pop, M. & Salzberg, S.L. Ultrafast and memory-efficient alignment of short DNA sequences to the human genome. Genome Biol 10, R25 (2009).

79. Li, H. et al. The Sequence Alignment/Map format and SAMtools. Bioinformatics 25, 2078–9 (2009).

80. Buenrostro, J.D., Wu, B., Chang, H.Y. & Greenleaf, W.J. ATAC-seq: A Method for Assaying Chromatin Accessibility Genome-Wide. Current protocols in molecular biology 109, 21.29.1–21.29.9 (2015).

81. Langmead, B. & Salzberg, S.L. Fast gapped-read alignment with Bowtie 2. Nat Methods 9, 357–9 (2012).

82. Jun, G. et al. Detecting and estimating contamination of human DNA samples in sequencing and array-based genotype data. Am J Hum Genet 91, 839–48 (2012).

83. Bates, D., Mächler, M., Bolker, B. & Walker, S. Fitting Linear Mixed-Effects Models Using lme4. Journal of Statistical Software 67, 1–48 (2015).

84. Kuznetsova, A., Brockhoff, P.B. & Christensen, R.H.B. lmerTest Package: Tests in Linear Mixed Effects Models. Journal of Statistical Software 82(2017).

85. Bocher, J. The BioVoxxel Image Processing and Analysis Toolbox. in EuBIAS-Conference (2015).

86. ThermoFisherScientificInc. Thermo Scientific Cellomics® Neuronal Profiling V4. in V4 Version (Thermo Fisher Scientific Inc., Pittsburgh, Pennsylvania 15219, USA).

87. Hartig, F. DHARMa: residual diagnostics for hierarchical (multi-level/mixed) regression models. (https://cran.r-project.org/, 2021).

88. Ongen, H., Buil, A., Brown, A.A., Dermitzakis, E.T. & Delaneau, O. Fast and efficient QTL mapper for thousands of molecular phenotypes. Bioinformatics 32, 1479–85 (2016).

89. Mohammadi, P., Castel, S.E., Brown, A.A. & Lappalainen, T. Quantifying the regulatory effect size of cis-acting genetic variation using allelic fold change. Genome Res 27, 1872–1884 (2017).

90. Kumasaka, N., Knights, A.J. & Gaffney, D.J. Fine-mapping cellular QTLs with RASQUAL and ATAC-seq. Nat Genet 48, 206–13 (2016).

91. Kuhn, R.M., Haussler, D. & Kent, W.J. The UCSC genome browser and associated tools. Brief Bioinform 14, 144–61 (2013).

92. Davis, J.R. et al. An Efficient Multiple-Testing Adjustment for eQTL Studies that Accounts for Linkage Disequilibrium between Variants. Am J Hum Genet 98, 216–24 (2016).

93. Hoffman, G.E. et al. CommonMind Consortium provides transcriptomic and epigenomic data for Schizophrenia and Bipolar Disorder. Sci Data 6, 180 (2019).

94. Krismer, K. et al. Transite: A Computational Motif-Based Analysis Platform That Identifies RNA-Binding Proteins Modulating Changes in Gene Expression. Cell Rep 32, 108064 (2020).

